# Oxidative stress causes a reversible decrease of deubiquitylases activity in old vertebrate brains

**DOI:** 10.1101/2025.08.15.670606

**Authors:** Amit Kumar Sahu, Alberto Minetti, Domenico Di Fraia, Antonio Marino, Patrick Rainer Winterhalter, Daniela Giustarini, Andreas Simm, Francesco Neri, Federico Galvagni, Christoph Gerhardt, Thorsten Pfirrmann, Alessandro Ori

**Affiliations:** Leibniz Institute on Aging - Fritz Lipmann Institute (FLI), Jena, Germany; Clinic for Heart Surgery (UMH), Martin-Luther-University Halle-Wittenberg, Halle (Saale), Germany; Department of Biotechnology, Chemistry and Pharmacy, University of Siena, Siena, Italy; Department of Life Sciences and Systems Biology, University of Turin, Torino, Italy; Health and Medical University, Potsdam, Germany

**Keywords:** Brain aging, Deubiquitylases (DUBs), Activity-based proteomics, Oxidative stress, Ubiquitin-Proteasome System (UPS), N-acetylcysteine ethyl ester (NACET)

## Abstract

The ubiquitin-proteasome system is essential for neuronal proteostasis, and its activity declines with age. How deubiquitylating enzymes (DUBs) are affected by aging in the vertebrate brain remains unclear. Here, we profiled cysteine protease DUBs using activity-based proteomics in aging mouse and killifish brains. Despite stable protein levels, we identified a subset of DUBs that progressively lose catalytic activity with age. We demonstrated that oxidative stress impairs DUB function through thiol oxidation and that antioxidant treatment restores their activity *in vitro* and *in vivo*. Further, inhibition of DUBs in human iPSC-derived neurons significantly recapitulated ubiquitylation changes observed in aged brains, and temporal analysis in mice revealed that DUB inhibition precedes proteasome decline in the brain during aging. Together, these findings indicate a redox-sensitive subset of DUBs that undergo an age-associated decline in activity and suggest that impaired deubiquitylation is an early, yet potentially reversible, driver of proteostasis decline in the aging brain.

## Introduction

Aging is characterized by widespread functional and molecular changes in the brain, including synaptic dysfunction, increased oxidative stress, and altered protein homeostasis ^1^. Among the cellular mechanisms governing proteostasis, the ubiquitin-proteasome system (UPS) plays a central role in signaling, stress responses, and protein degradation by attaching ubiquitin to lysine residues of specific target proteins ^2^. The decline in proteasome activity and alterations in the ubiquitylome have been well characterized in the context of aging in both invertebrates and vertebrates ^2–5^. Within the UPS, ubiquitin ligases and deubiquitylases (DUBs) act antagonistically to modulate protein fate and signaling pathways dynamically.

Among these enzymes, DUBs constitute a diverse family of more than 100 enzymes identified in vertebrates, broadly classified into cysteine proteases and zinc-dependent metalloproteases ^6^. DUBs are central regulators of the cell cycle, immune signaling, and control multiple aspects of neuronal function, including mitochondrial quality control, synaptic plasticity, and stress responses ^6^. Altering DUB activity has been linked to lifespan in nematodes ^4^, and dysregulation of specific DUBs in humans leads to several neurodegenerative diseases, such as spinocerebellar ataxia and Parkinson’s disease ^7–10^. Emerging evidence has implicated the tight regulation of DUB activity by protein binding and post-translational modifications, including the redox regulation of the cysteine protease DUBs ^6,11^. However, a systematic understanding of how DUB function is altered in the aging brain, the mechanisms driving these changes, and the consequences of altered DUB activity at the molecular level are still lacking.

To fill this gap, we profiled the activity of cysteine-dependent DUBs in mouse (*Mus musculus*) and killifish (*Nothobranchius furzeri*) brains across different age groups. Using activity-based probes coupled with mass spectrometry and biochemical validation, we found that the average DUB activity is reduced by approximately 40% in old brains due to increased cysteine oxidation, and it can be restored by antioxidant treatments both *in vitro* and *in vivo*. By employing human-induced pluripotent stem cell (iPSC)-derived neurons (iNeurons), we demonstrate that DUB inhibition can recapitulate a significant fraction of age-related ubiquitylated signatures identified in the mouse brains. Importantly, we show that decreased DUB activity temporally precedes impairment of proteasome activity and accumulation of ubiquitylated proteins during brain aging. Together, our findings revealed a conserved susceptibility of DUBs to redox imbalance and provided insights into impaired deubiquitylation as a contributing factor to age-associated proteostasis decline.

## Results

### Global decrease in enzymatically active deubiquitylases in aging vertebrate brains

To understand how the activity of DUBs changes during the aging process in mouse brains, we performed a DUBs activity assay by utilizing a fluorescent deubiquitylase substrate, as previously used in ^12^. We observed a decrease in the global DUB activity in old (30 months) mouse brain lysates compared to young (3 months) ones, which cannot be explained by changes in DUB protein abundance (proteomics data from ^5^) (Fig. 1A, Fig. S1A). To gain deeper insight into the activity of individual DUBs, we used activity-based probes to enrich cysteine protease DUBs and their identification using a mass spectrometry (MS) approach. To maximize the capture of individual DUBs, we pooled probes from three different classes that included a mono-ubiquitin recognition element specific for DUBs, a biotin-tag for DUBs’ pulldown, and either a propargylamide (PA), a vinylmethylester (VME), or a vinylsulfone (VS) electrophilic warhead to covalently interact with the nucleophilic cysteine residue in the active sites of DUBs ^13–15^ (Fig. 1B). Hereafter, we referred to the pool of activity-based probes as DUB probes. As a negative control, we pre-treated brain lysates with N-ethylmaleimide (NEM) that can irreversibly alkylate any free sulfhydryl group, such that DUB probes will fail to bind and enrich DUBs in the pulldown assay (Fig. 1B, Fig. S1B). The Principal Component Analysis (PCA) of enriched proteins depicted a clear separation based on age groups and DMSO (vehicle) or NEM treatment (Fig. 1C). Next, we defined the quantity of DUBs being enzymatically active as the abundance of DUBs enriched over the NEM control. This revealed the significant enrichment (|average Log_2_ FC| > 0.58; Qvalue < 0.05) of 44 and 45 active DUBs in young and old brains, respectively. In addition, despite implementing stringent washing steps during our enrichment process, we still observed the co-enrichment of DUB substrates and interactors due to their physical association with DUBs, as reported previously by ^16,17^ (Fig. 1D, Fig. S1C). In line with the fluorescent DUB substrate-based assay, we observed an overall decrease in the activity of DUBs in old brains compared to young ones (Fig. 1E and 1F). Among significantly affected DUBs, we identified UCHL1 (a neuron-specific DUB) and YOD1 (a DUB broadly expressed across brain cell types) that have been linked to neurodegenerative diseases ^18–23^ (Fig. S1D). On the other hand, the activity of some other neurodegeneration-associated DUBs, such as CYLD and USP9X ^24–26^, was not altered by aging (Fig. S1D). Of note, we also observed the enrichment of metalloprotease DUBs, such as PSMD7, PSMD14, and COPS5, among others, due to their physical interaction with cysteine-dependent DUBs. However, their enrichment showed no significant difference between young and old ^17^ (Fig. S1E).

**Figure 1:**
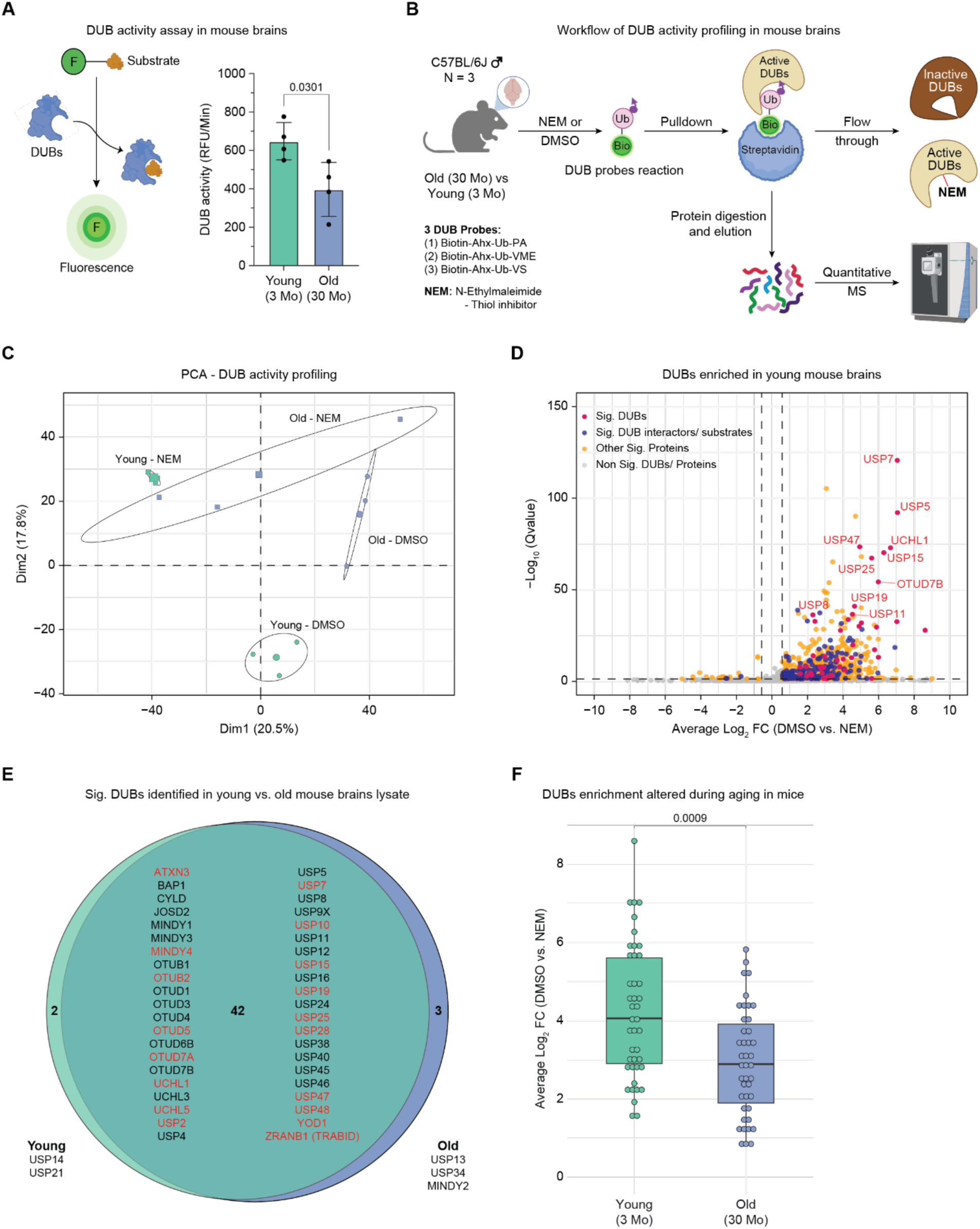
DUB activity decreases during mouse brain aging. (A) Left: Schematic illustrating the mechanism of fluorescent substrate-based DUB activity measurement. Right: DUB activity in brain lysates from young (3 months) and old (30 months) C57BL/6J male mice (N = 4 biological replicates; unpaired t-test with Welch’s correction; RFU = Relative Fluorescence Units; see Supplementary Table 1). (B) Experimental workflow for enrichment and characterization of active DUBs using activity-based probes in young and old C57BL/6J male mouse brains (N = 3 biological replicates; Ub = mono-ubiquitin molecule; Bio = biotin tag; arrowheads indicate probe warheads: propargylamide [PA], vinylmethylester [VME], or vinylsulfone [VS]). (C) Principal component analysis (PCA) of proteins enriched by DUB probes from mouse brains. Ellipses highlight sample groups (young vs. old brains treated with either DMSO or NEM) and represent 95% confidence intervals. The percent variance explained by each principal component is indicated. (D) Volcano plot of proteins enriched from young mouse brains. Vivid pink dots indicate active DUBs; deep indigo dots mark known DUB substrates and interactors from ^16,17^; vivid yellow dots denote other significantly co-enriched proteins; light grey dots indicate non-significant proteins. Seven non-DUB proteins were excluded from the plot for clarity (Qvalues from Spectronaut differential abundance analysis). Dashed lines indicate the threshold used to select differentially enriched proteins (|average Log2 FC| > 0.58; Qvalue < 0.05). (E) Venn diagram showing the overlap of active DUBs enriched from young and old brain lysates. DUBs in red significantly differ between age groups (normalized to respective NEM-treated samples; |average Log2 FC| > 0.58; Qvalue < 0.05; see Supplementary Table 2). (F) Boxplot of individual active DUB enrichment (y-axis) in young and old mouse brains (x-axis). Data includes 42 DUBs commonly detected in both groups (|average Log2 FC| > 0.58; Qvalue < 0.05; Wilcoxon rank-sum test). Related to Supplementary Tables 1 and 2.

**Supplementary Figure 1:**
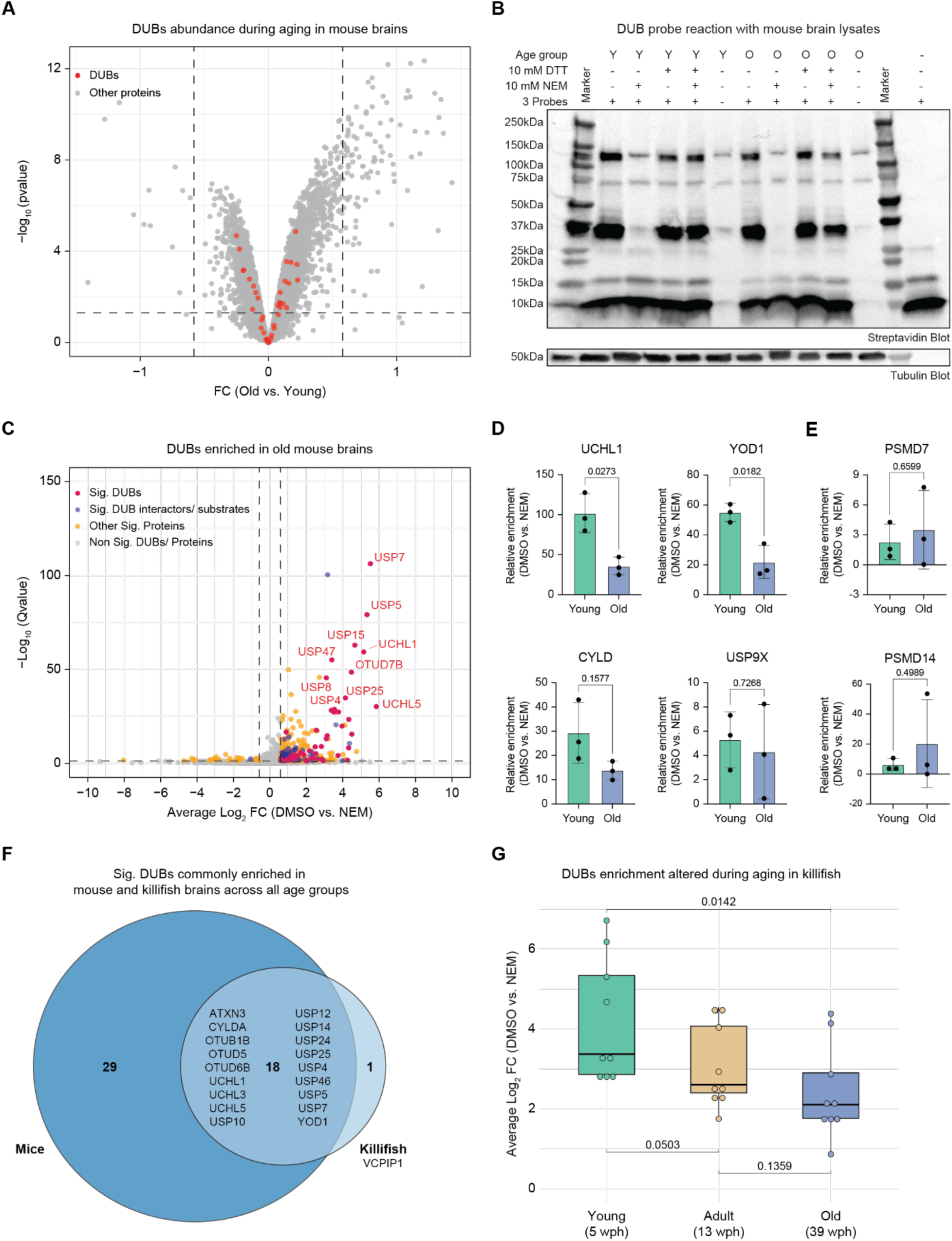
DUB activity in mouse and killifish brains during aging. (A) Volcano plot showing the abundance of DUB proteins in old (33 months) vs. young (3 months) C57BL/6J male mouse brains from ^5^. The x-axis was constrained from -1 to 1 for visualisation, owing to removing 12 significant non-DUB proteins (N = 5 biological replicates; |average Log2 FC| > 0.58; Qvalue < 0.05). (B) Immunoblot validating the reaction of DUB probes with young and old mouse brain lysates pre-treated either with DMSO, NEM, or DTT. Immunoblots were repeated for all 3 biological replicates used in DUB activity profiling in mouse brains (N = 3 biological replicates). (C) Volcano plot of proteins enriched from old mouse brains. Vivid pink dots indicate active DUBs; deep indigo dots mark known DUB substrates and interactors from ^16,17^; vivid yellow dots denote other significantly co-enriched proteins; light grey dots indicate non-significant proteins. Seven non-DUB proteins were excluded from the plot for clarity (Qvalues from Spectronaut differential abundance analysis). Dashed lines indicate the threshold used to select differentially enriched proteins (|average Log2 FC| > 0.58; Qvalue < 0.05). (D) Bar plot depicting examples of enrichment of cysteine protease DUBs (N = 3 biological replicates; unpaired t-test with Welch’s correction). (E) Bar plot depicting examples of enrichment of metalloprotease DUBs (N = 3 biological replicates; unpaired t-test with Welch’s correction). (F) Venn diagram showing the overlap of active DUBs enriched in all age groups from mouse and killifish. Of note, VCPIP1 is enriched in young and old mouse brain lysates treated with DTT (normalized to respective NEM-treated samples; |average Log2 FC| > 0.58; Qvalue < 0.05). (G) Boxplot of individual active DUB enrichment (y-axis) in young (5wph), adult (13wph), and old (39wph) killifish brains (x-axis). Data includes 9 DUBs commonly detected in all age groups (|average Log2 FC| > 0.58; Qvalue < 0.05; Wilcoxon rank-sum test; wph = weeks post hatching). Related to Supplementary Table 2.

We have previously shown that changes in protein ubiquitylation and decreased proteasome activity are conserved aging signatures in killifish and mice ^3,5,27^. To understand whether the decrease in DUB activity during brain aging is also a conserved phenotype across vertebrate species, we repeated our DUB probe-based activity profiling with brain lysates of killifish of different age groups. 19 DUBs showed significant enrichment relative to the NEM control across all age groups in killifish, of which 18 are consistently identified in young and old mouse brain lysates (Fig. S1F, Supplementary Table 2). Consistent with mouse data, we observed a decrease in global DUB activity in old killifish brains compared to the young ones (Fig. S1G). Together, these results show that the activities of DUBs are reduced in both old mouse and killifish brains compared to young animals. The decrease of DUB activity is largely independent of protein abundance (Fig. S1A), suggesting the involvement of post-translational mechanisms influencing DUB activity in old brains.

### Oxidative stress-mediated thiol oxidation contributes to the reduced deubiquitylase activity in old brains

Elevated levels of reactive oxygen species (ROS), along with chronically activated antioxidant defense mechanisms that may become insufficient over time, have been implicated in aging and age-associated diseases, contributing to impaired protein homeostasis and reduced enzymatic function ^1,28^. Consistently, in proteome data from aged mouse brains ^5^, we observed increased abundance of antioxidant enzymes such as Apolipoprotein D (APOD), Selenoprotein K (SELENOK), and Nuclear factor erythroid 2-related factor 2 (NRF2) target proteins, along with a decrease in abundance of protective factors like Serine protease inhibitor A3K (SERPINA3K), Reactive oxygen species modulator 1 (ROMO1), and Haptoglobin (HP) (Fig. S2A). Since many DUBs rely on the thiol of cysteine residues at their active site to cleave ubiquitin molecules from substrates or another ubiquitin molecule, we hypothesized that oxidative stress-mediated thiol oxidation in the brains of old mice might explain the reduced enrichment of active DUBs in our experiments ^29^. To test this hypothesis, we quantified the concentration of reduced form of thiols (-SH) in young and old mouse brains’ lysate using Ellman’s reagent, 5,5’-dithio-bis-(2-nitrobenzoic acid) (DTNB), which is also used to study cysteine oxidation ^30,31^. We observed a reduction in thiol groups in old brain lysates compared to young ones in two independent experiments, suggesting that thiol oxidation increases with age (Fig. 2A). Further, to understand whether the DUB activity depends on the concentration of reduced thiol groups in cysteines, we performed a linear regression analysis. We observed a statistically significant linear relationship (P value = 0.042, R² = 0.684), confirming that thiol concentration (or, in other words, cysteine’s oxidation) correlates significantly with DUB activity in the same set of samples (Fig. S2B).

**Figure 2:**
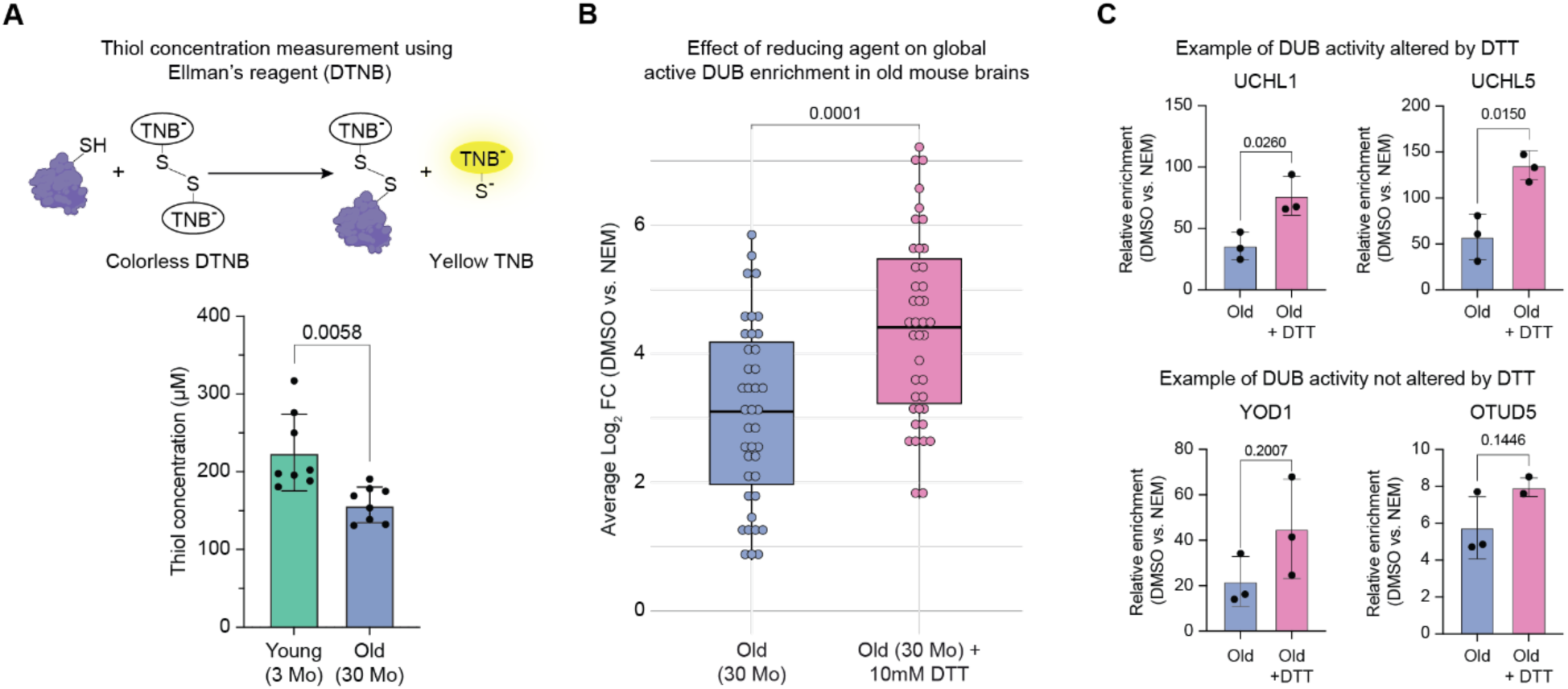
Increased thiol oxidation leads to a reversible decrease in DUB activity during mouse brain aging. (A) Upper: Schematic illustrating the reaction of Ellman’s reagent (DTNB) with reduced thiol groups, producing the yellow TNB product measurable at 412 nm via spectrophotometry. Lower: Reduced thiol concentrations in young (3 months) and old (30 months) C57BL/6J male mouse brains (data pooled from two independent experiments with N = 3 and N = 5 biological replicates; unpaired t-test with Welch’s correction). (B) Boxplot of individual active DUB enrichment (y-axis) in old mouse brains with or without 10 mM DTT treatment (x-axis). Data includes 40 DUBs commonly detected in old brain lysates ± DTT (N = 3 biological replicates; |average Log2 FC| > 0.58; Qvalue < 0.05; Wilcoxon rank-sum test). (C) Bar plot depicting examples of cysteine protease DUBs enrichment in old brain lysates treated with (upper) or without 10 mM DTT (lower) (N = 3 biological replicates; unpaired t-test with Welch’s correction). Related to Supplementary Tables 1 and 2.

**Supplementary Figure 2:**
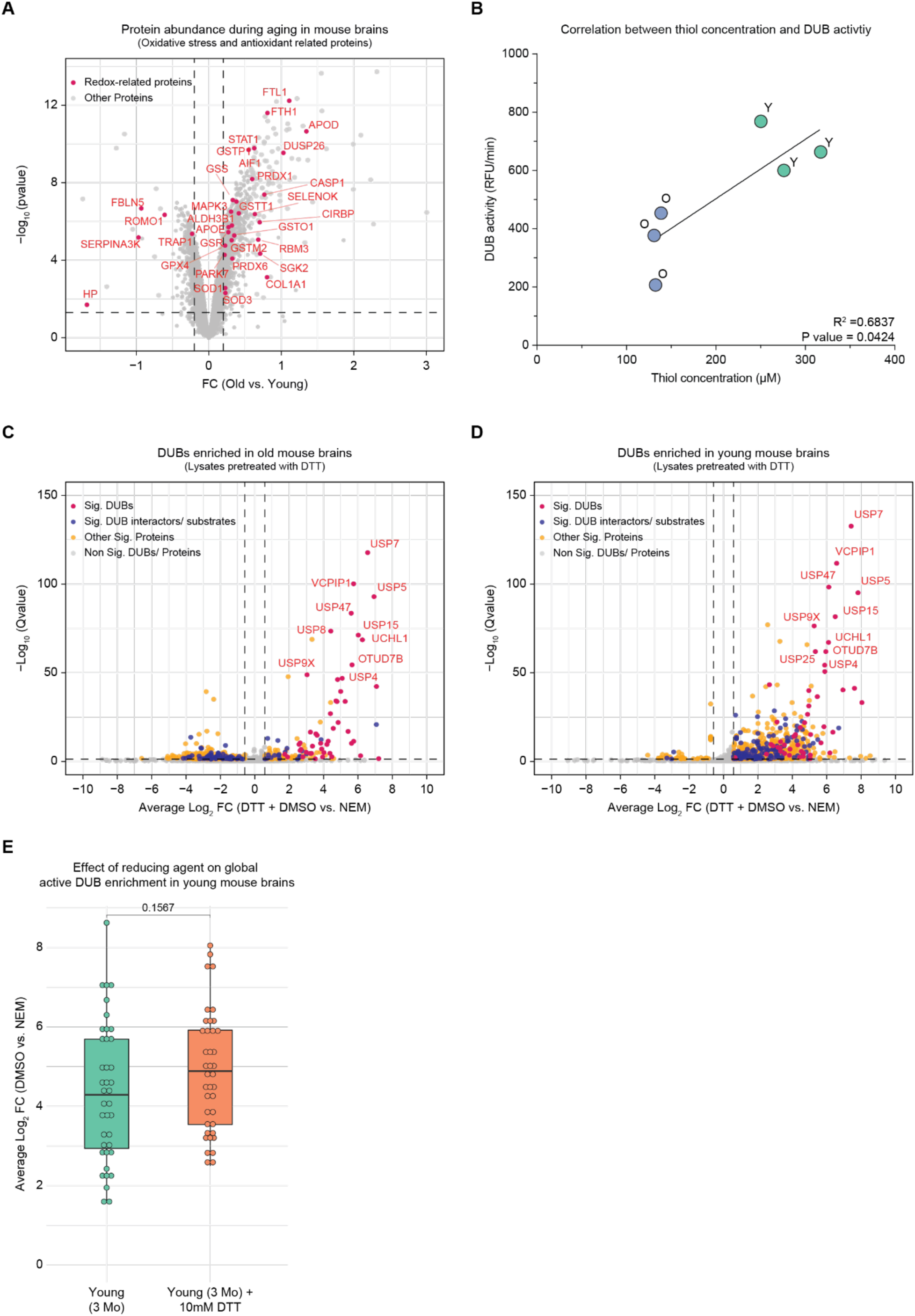
Oxidative stress-mediated reversible cysteine oxidation modulates DUB activity during brain aging. (A) Volcano plot showing the abundance of redox-related proteins in old (33 months) vs. young (3 months) C57BL/6J male mouse brains from ^5^. Red dots indicate redox-related proteins (N = 5 biological replicates; |average Log₂ FC| > 0.2; Qvalue < 0.05; pvalues from Spectronaut differential abundance analysis). Dashed lines indicate the threshold used to select differentially enriched proteins. (B) Linear regression analysis between thiol concentration (μM, x-axis) and DUB activity (RFU/min, y-axis). The solid line represents the best-fit regression line (R² = 0.6837; P = 0.0424). (C) Volcano plot of proteins enriched from old mouse brain lysates treated with 10 mM DTT. Seven non-DUB proteins, one DUB interactor, and three non-significant proteins were excluded for clarity. (D) Volcano plot of proteins enriched from young mouse brain lysates treated with 10 mM DTT. Three non-DUB proteins and three non-significant proteins were excluded for clarity. Vivid pink dots indicate active DUBs; deep indigo dots mark known DUB substrates and interactors ^16,17^; vivid yellow dots denote other significantly co-enriched proteins; light grey dots represent non-significant proteins (Qvalues from Spectronaut differential abundance analysis). Dashed lines indicate the threshold used to select differentially enriched proteins (|average Log2 FC| > 0.58; Qvalue < 0.05). (E) Boxplot of individual active DUB enrichment (y-axis) in young mouse brains with or without 10 mM DTT treatment (x-axis). Data includes 39 DUBs commonly detected in young brain lysates ± DTT (N = 3 biological replicates; |average Log₂ FC| > 0.58; Qvalue < 0.05; Wilcoxon rank-sum test). Related to Supplementary Tables 1 and 2.

Next, we asked whether we could reverse the effect of oxidation on the activity of DUBs in old brain lysates. We treated brain lysates from old mice with dithiothreitol (DTT), a reducing reagent previously shown to enhance DUB activity *in vitro ^11^*, followed by DMSO or NEM and DUB probe-based enrichment, as in (Fig. 1B). We observed a significant increase in the overall enrichment of active DUBs in old brain lysates treated with DTT compared to the non-treated lysate (Fig. 2B, Fig. S2C). Further, by performing the relative enrichment analysis of the protein quantity, we found an enhancement of individual DUB activity, such as that of UCHL1 and UCHL5, in DTT-treated old brain lysates. However, DTT treatment didn’t significantly alter the activity of other DUBs, e.g., YOD1 and OTUD5 (Fig. 2C). On the other hand, DTT did not impart significant changes in the enrichment of active DUBs in young brain lysates (Fig. S2D-E). These observations support that reduced DUB activity in old mouse brains might derive from increased cysteine oxidation.

### DUB inhibition partially recapitulates age-related ubiquitylation signatures

After observing decreased DUB activity (this study) and increasing ubiquitylated protein levels ^5^ during brain aging, we next sought to understand the effect of DUB inhibition on the ubiquitylated proteome. We acutely inhibited DUBs using a broad-spectrum DUB inhibitor, PR619 ^32^, in 14 days post-differentiated human iPSC-derived neurons ^33^ (Fig. 3A). We chose this model as it has been utilized previously to investigate proteome-wide protein ubiquitylation ^5,34,35^ and to study human age-associated neurodegenerative diseases ^36^. We chose a PR619 concentration and treatment duration that does not impart cellular toxicity (Fig. 3A, Fig. S3A) and minimizes off-target effects ^37^ while being sufficient to inhibit DUBs (Fig. S3B). Under these conditions, PR619 treatment does not directly affect proteasome activity (Fig. S3C). In parallel, we also inhibited proteasome function using bortezomib as a control (Fig. 3A, Fig. S3B-C). We observed the expected increase in the total ubiquitylated protein levels (Fig. S3D). Subsequently, we enriched ubiquitylated peptides using a well-established antibody-based approach specific to the lysine di-GLY (K-*ε*-GG) remnant motif ^38^ (Fig. 3A). Although this method leads to the enrichment of other modifications, such as ISGylation and NEDDylation, more than 95% of K-*ε*-GG-modified sites are contributed by ubiquitylation ^39^. Hence, from this point forward, we refer to K-*ε*-GG-modified sites as ubiquitylated. In parallel, we measured proteome changes of treated-iNeurons using DIA-based MS (Fig. 3A). We noted a robust increase in the number of identified ubiquitylated peptides upon PR619 and bortezomib treatment (>14,000 Ub sites) compared to the DMSO (>6,000 Ub sites). In contrast, the number of total proteins quantified remained similar (∼7000 proteins) (Fig. 3B, Fig. S3E).

**Figure 3:**
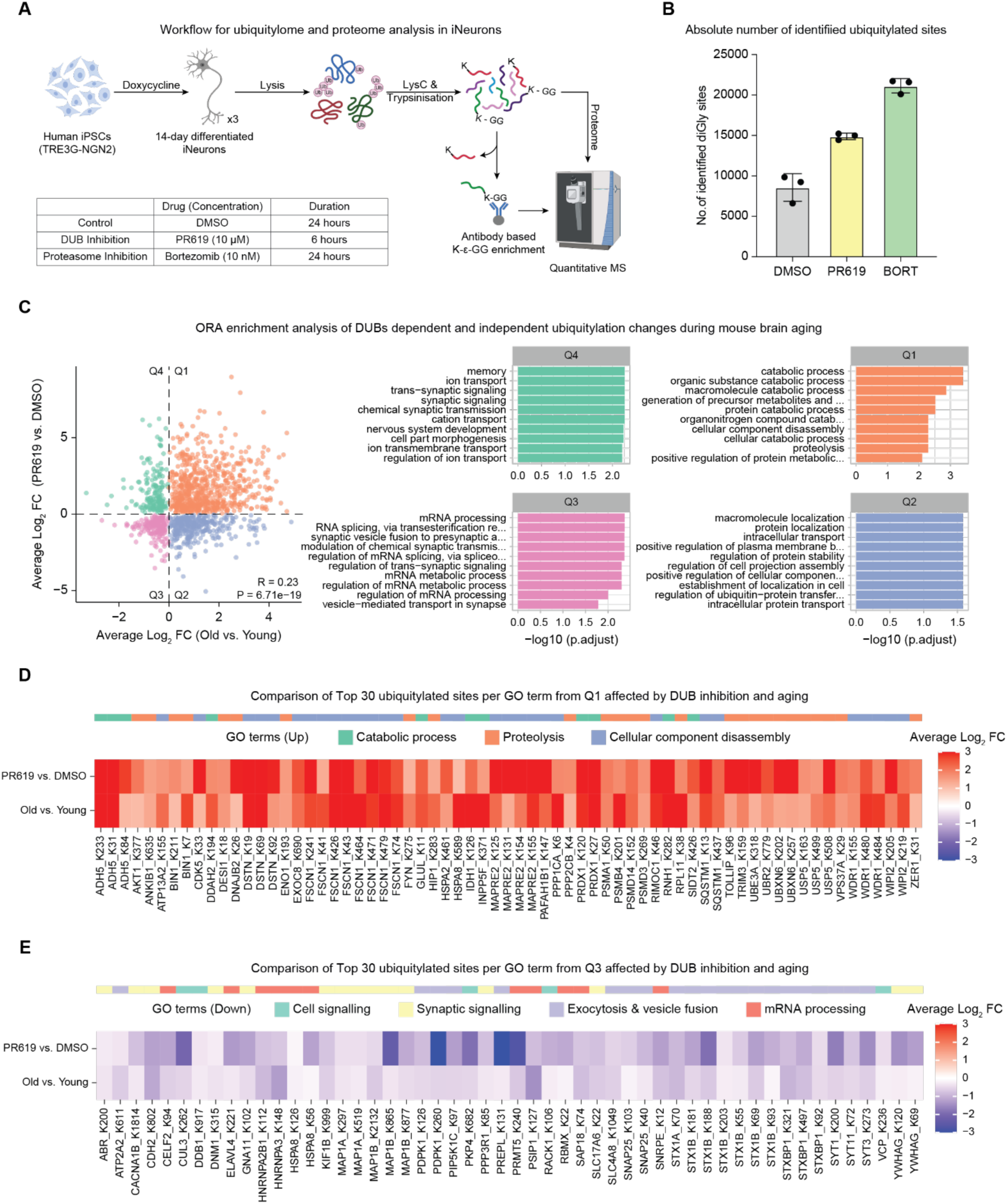
DUB inhibition contributes to age-related protein ubiquitylation signatures. (A) Schematic illustrating the K-ε-GG antibody-mediated ubiquitylated peptide enrichment and total proteome analysis of iPSC-derived iNeurons treated either with DMSO, 10 µM PR619, or 10 nM Bortezomib (N = 3 biological replicates). (B) Bar plot depicting the number of identified ubiquitylated sites in iNeurons treated with different drugs (error bars represent the standard deviation from the mean). (C) Left: Scatter plot comparing differentially enriched ubiquitylated sites between mouse aging (old vs. young, x-axis, from ^5^) and DUB-inhibited iNeurons (PR619 vs. DMSO, y-axis). Right: Quadrant-based Over Representation Analysis (ORA) of the top 10 biological processes. Data includes ubiquitylated site changes with adj.pvals < 0.05 (for mouse) and Qvalue < 0.05 (for iNeurons). (D) Heatmap representing the comparison of average Log2 FC of the top 30 enriched ubiquitylated sites per GO term from Quadrant 1 (Q1) in (C) between mouse aging and DUB-inhibited iNeurons. (E) Heatmap representing the comparison of average Log2 FC of the top 30 enriched ubiquitylated sites per GO term from Quadrant 3 (Q3) in (C) between mouse aging and DUB inhibited iNeurons (for both heatmaps, only differentially enriched sites with adj.pvals < 0.05 (for mouse) and Qvalue < 0.05 (for iNeurons) were used). Related to Supplementary Table 3.

**Supplementary Figure 3:**
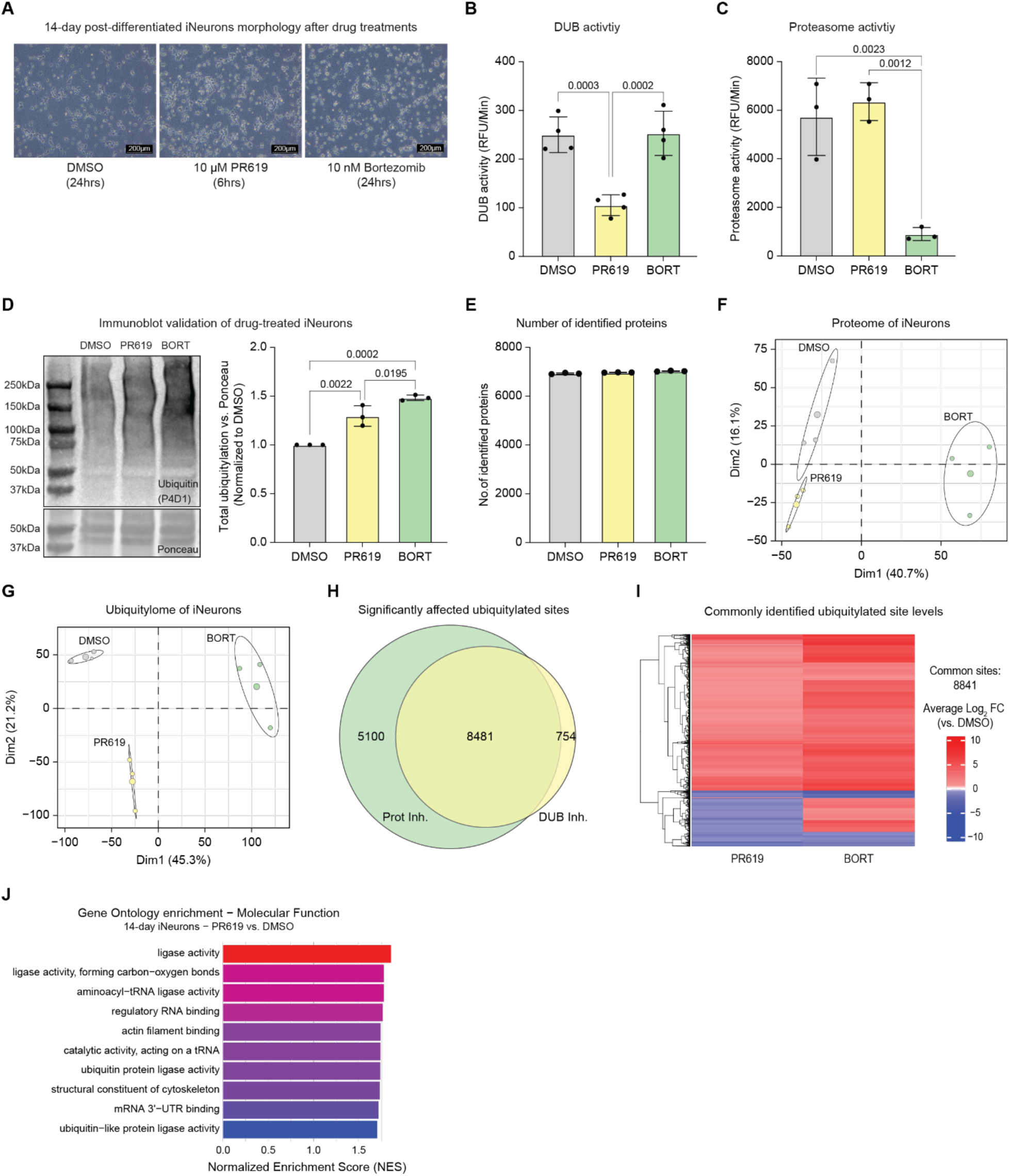
Impact of DUB inhibition on proteome and ubiquitylome in iNeurons. (A) Representative images showing the morphology of 14 days post-differentiated iNeurons after treatment with DMSO, 10 µM PR619, or 10 nM Bortezomib at the indicated time points (magnification = 10X; scale = 200 µm; repeated across N = 3 biological replicates). (B) DUB activity in iNeurons. (C) Proteasome activity in iNeurons following drug treatment. In both panels, N = 4 biological replicates; one-way ANOVA; RFU = Relative Fluorescence Units. (D) Left: Immunoblot confirms increased levels of total ubiquitylated proteins in iNeurons after PR619 and Bortezomib treatments. Total protein levels were assessed using Ponceau staining. Immunoblots were repeated for all 3 biological replicates used in ubiquitylated site enrichment with K-ε-GG antibody and proteome analysis. Right: Quantification of total ubiquitylation normalized to total protein levels across treatment conditions (N = 3 biological replicates; one-way ANOVA). (E) Bar plot showing the number of identified proteins in iNeurons treated with different drugs (error bars represent the standard deviation from the mean). (F) PCA of proteome changes in iNeurons. (G) PCA of ubiquitylome changes in iNeurons. Ellipses highlight drug treatment groups in both PCA plots and represent 95% confidence intervals. The percent variance explained by each principal component is indicated. (H) Venn diagram showing the overlap in the number of differentially enriched ubiquitylated sites between DUB inhibition (PR619 vs. DMSO) and proteasome inhibition (Bortezomib vs. DMSO) (|average Log2 FC| > 0.58; Qvalue < 0.05). (I) Heatmap displaying hierarchical clustering (Euclidean distance) of 8,481 ubiquitylated sites commonly identified in DUB and proteasome-inhibited iNeurons. The column represents treatment conditions, and rows show average log2 FC intensity of each ubiquitylated site compared to DMSO (|average Log2 FC| > 0.58; Qvalue < 0.05). (J) Gene Set Enrichment Analysis (GSEA) of ubiquitylated sites altered in DUB-inhibited iNeurons (PR619 vs. DMSO). The top 10 Gene Ontology (GO) terms associated with increased ubiquitylation, ranked by NES, are shown. Related to Supplementary Tables 1 and 3.

PCA based on global proteome data revealed that DUB inhibition by PR619 had a reduced impact on protein abundance compared to proteasome inhibition by bortezomib (Fig. S3F). On the other hand, both DUB and proteasome inhibition strongly affected the ubiquitylome compared to the vehicle control (Fig. S3G). Euclidean distance-based hierarchical clustering of 8481 cross-quantified ubiquitylated peptides revealed a more pronounced effect of bortezomib on ubiquitylated sites than PR619, as expected. Most of the commonly affected sites exhibited consistent changes (6251 sites showing increased and 824 decreased ubiquitylation, Fig. S3H-I). These data show that DUB inhibition leads to changes in protein ubiquitylation that are less pronounced but largely overlapping with proteasome inhibition. To gain a deeper understanding of the molecular function that might be impacted by DUB inhibition, we performed gene set enrichment analysis (GSEA) using proteins that showed altered ubiquitylation. We observed that proteins belonging to ligase activity, actin filament binding, cytoskeleton, and mRNA binding were among the top 10 terms associated with increased ubiquitylation following DUB inhibition (Fig. S3J).

To assess whether the ubiquitylated sites altered by DUB inhibition resemble those affected by aging, we compared significantly altered ubiquitylated sites in PR619-treated iNeurons with those observed in aging mouse brains ^5^ (Fig. 3C). Pearson correlation analysis revealed that DUB inhibition significantly and partially recapitulated aspects of age-induced ubiquitylated signatures (R = 0.22, P = 4.4e-15), suggesting that altered DUB activity may contribute to age-related molecular changes (Fig. 3C). To explore this overlap further, we analyzed GO terms enriched among ubiquitylated sites affected by both DUB inhibition and aging (Fig. 3C). Terms related to catabolic processes and proteolysis were among the top 10 enriched categories with increased ubiquitylation in both aging and following DUB inhibition. For example, increased ubiquitylation of proteins belonging to catabolic processes such as alcohol dehydrogenase 5 (ADH5), an enzyme involved in glutathione-dependent formaldehyde detoxification pathway and maintaining redox balance in cells ^40^ was observed under both aging and following DUB inhibition (Fig. 3D). Similarly, increased ubiquitylation of proteasome-associated subunits like PSMA1 and PSMB4, and autophagy-related proteins such as sequestosome-1 (SQSTM1) was detected across multiple sites (Fig. 3D). On the other hand, proteins involved in synaptic signaling, exocytosis and vesicle fusion, such as syntaxin-1B (STX1B) and its binding protein (STXBP1) showed reduced ubiquitylation during aging and following DUB inhibition (Fig. 3E). Together, these results demonstrate that the reduction of DUB activity may contribute to some of the age-associated ubiquitylation changes observed in the brains of aged mice.

### Decline in DUB activity precedes proteasome impairment during aging in mouse brains

Since both DUB and proteasome inhibition ^5^ can independently contribute to aging-like signatures, and emerging evidence suggests they may influence each other’s function ^41–45^, we next investigated which of these molecular events occurs earlier during the aging process. To investigate this, we performed a temporal study analyzing DUB activity and proteasome activity, along with measuring the concentration of the reduced form of thiol (-SH) and total ubiquitylated protein levels in the brains of mice of different ages (Fig. 4A). We observed a stable thiol concentration until 12 months and then a significant decrease beginning from 18 months of age (Fig. 4B). In accordance with changes in thiol concentration, we observed a decline in DUB activity post-18 months of age compared to younger ages (Fig. 4C). On the other hand, we noticed a gradual accumulation of ubiquitylated proteins in the brain with age, mirroring the decrease in the DUB activity (Fig. 4D, Fig. S4A). Interestingly, we found a minor fluctuation in the activity of the proteasome across ages and significantly reduced activity emerging only at 30 months (Fig. 4E). A similar decline in DUB (Fig. S1G) and proteasome activity ^3^ has been observed in middle-aged killifish brains (12-13 wph). To assess whether changes in DUB and proteasome activity are associated with declining thiol concentrations during aging, we performed correlation analyses. We found that younger brains with higher thiol levels exhibit increased DUB and proteasome activity, while aged brains with lower thiol concentrations show reduced activity. Notably, the correlation with thiol concentration was stronger for DUB activity than for proteasome activity (Fig. S4B).

**Figure 4:**
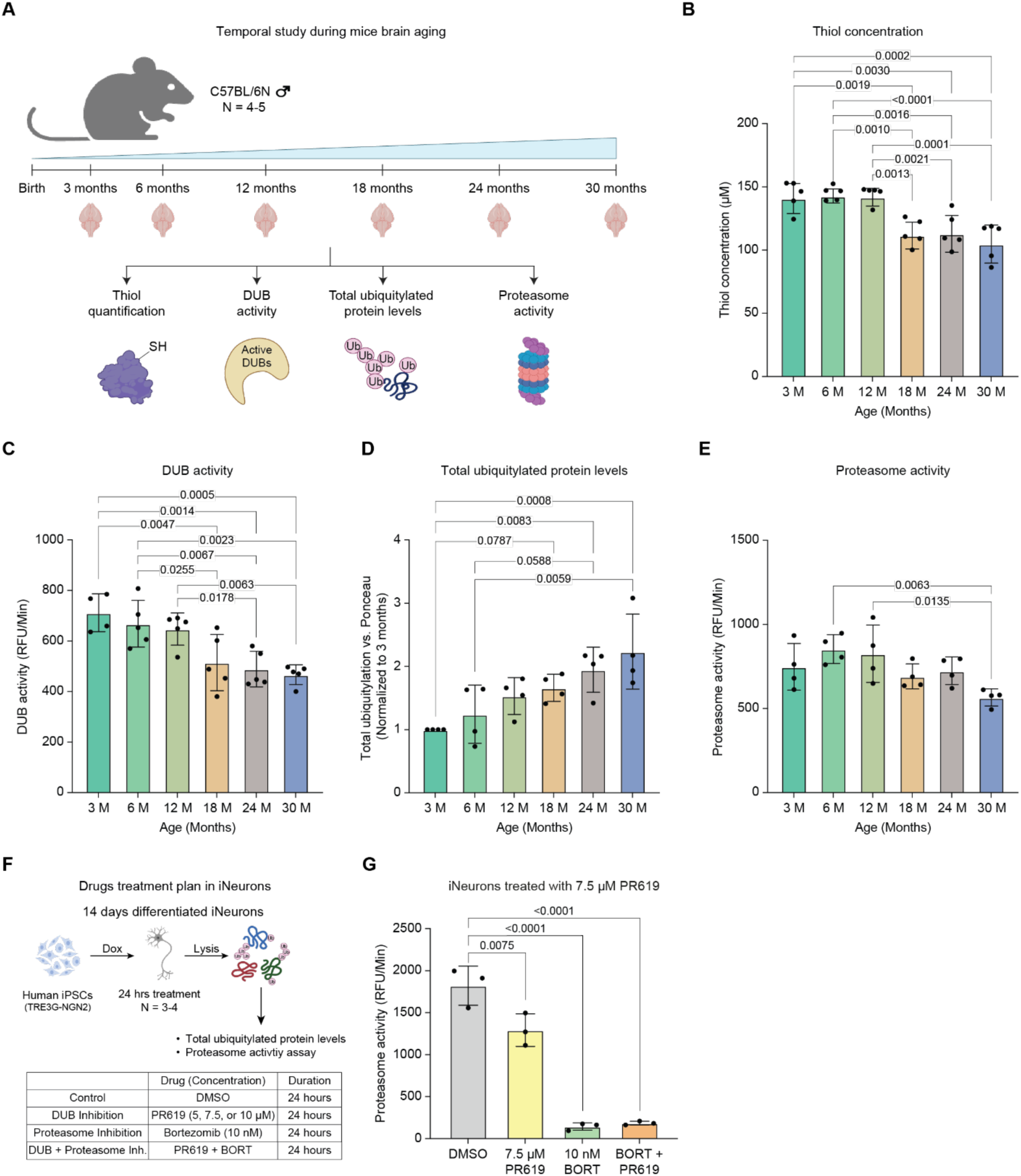
Temporal study of molecular changes occurring during mouse brain aging. (A) Schematic illustrating the temporal study of molecular changes during aging in C57BL/6N male mouse brains (N = 4-5 biological replicates). (B) Reduced thiol concentrations in the mouse brains of varying age groups (N = 5). (C) DUB activity (N = 4-5). (D) Total ubiquitylated protein levels (N = 4). (E) Proteasome activity (N = 4). (F) Schematic of chronic DUB inhibition in iNeurons using different PR619 concentrations and 10 nM Bortezomib for 24 hours (N = 3-4). (G) Proteasome activity of iNeurons treated with 7.5 µM PR619, 10 nM Bortezomib, and a combination of both drugs for 24 hours (N = 3). N represents biological replicates in all panels, and one-way ANOVA was used for analysis. RFU = Relative Fluorescence Units; Dox = Doxycycline. Related to Supplementary Table 1.

**Supplementary Figure 4:**
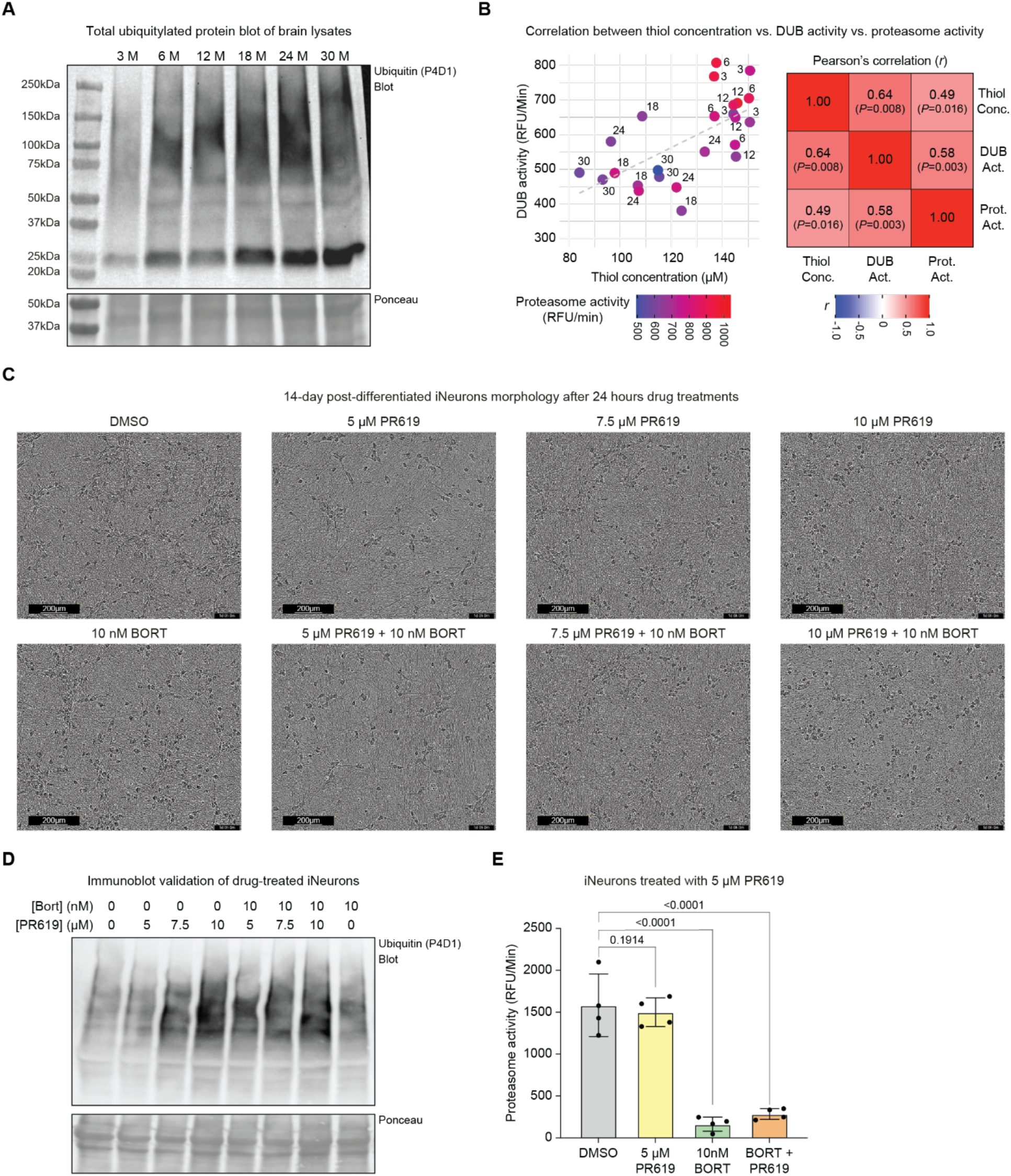
Effects of DUB inhibition on ubiquitylated protein levels and proteasome activity in mouse brain and iNeurons. (A) Immunoblot analysis of total ubiquitylated protein levels in the mouse brains across different age groups. Total protein levels were assessed using Ponceau staining. Immunoblots were repeated for all 5 biological replicates; technical outliers were excluded. (B) Left: Scatter plot showing the relationship between thiol concentration (X-axis), DUB activity (Y-axis), and proteasome activity (represented by bubble color) in mouse brains of the indicated age groups. Right: Pearson’s correlation analysis of the molecular changes associated with brain aging. (C) Representative images showing the morphology of 14 days post-differentiated iNeurons after 24 hours treatment with varying concentrations of PR619, with or without 10 nM Bortezomib (magnification = 10X; scale = 200 µm; repeated across N = 3-4 biological replicates). (D) Immunoblot validation of total ubiquitylated protein levels in iNeurons following treatment with different concentrations of PR619 and 10 nM Bortezomib. Total protein levels were assessed using Ponceau staining. (E) Proteasome activity of iNeurons treated for 24 hours with 5 µM PR619, 10 nM Bortezomib, or their combination (N = 4 biological replicates; one-way ANOVA; RFU = Relative Fluorescence Units). Related to Supplementary Table 1.

We next asked whether changes in DUB activity underlie the reduction in proteasome activity during aging. Earlier, we observed that acute inhibition of DUBs in iNeurons using 10 µM PR619 for 6 hours didn’t alter proteasome activity (Fig. S3C). Thus, we hypothesized that chronic inhibition of DUBs could impact the proteasome. To test this, we inhibited DUBs in iNeurons using PR619 for 24 hours at three different concentrations and bortezomib as a positive control (Fig. 4F). PR619 showed toxicity at 10 µM after 24 hours, while 5 µM and 7.5 µM were non-toxic, even when combined with bortezomib. While 5 µM had no effect, 7.5 µM PR619 increased ubiquitylated proteins and significantly reduced proteasome activity (Fig. 4G, Fig. S4C-E). Together, these results show that the decrease of DUB activity temporally precedes proteasome inhibition in the aging mouse brain, and that chronic impairment of DUBs in neurons could partially inhibit proteasome activity.

### NACET treatment rescues age-associated molecular changes in aged mouse brains

Building on our finding that DUB impairment (Fig. 1A and 1F) during brain aging is linked to oxidative stress and can be reversed (Fig. 2A and 2B), we tested whether antioxidant treatment could restore DUB function in aged animals. N-acetylcysteine ethyl ester (NACET) is a potent, orally bioavailable antioxidant known to cross blood-brain and blood-retinal barriers. Once inside cells, NACET is rapidly de-esterified to N-acetylcysteine (NAC), which is then gradually deacetylated, providing a sustained supply of cysteine ^46,47^ (Realini et al., accompanying manuscript). As cysteine is a precursor to glutathione (GSH), a major cellular antioxidant, we hypothesized that NACET could restore redox balance and DUB activity. To test this, aged mice (22 - 24 months) were treated with NACET in drinking water for 12 days (Fig. 5A). NACET treatment significantly increased the pool of reduced thiol groups in aged brains compared to vehicle controls (Fig. 5B), which was accompanied by restoration of DUB activity (Fig. 5C). Given that age-related DUB decline contributes to ubiquitylated protein accumulation and proteasome dysfunction, we next assessed whether NACET could rescue these molecular changes. While total ubiquitylated protein levels showed a trend toward decreased accumulation, K48-linked polyubiquitylated protein levels were significantly reduced in NACET-treated brains (Fig. 5D, Fig. S5A). Consistently, proteasome activity was enhanced following NACET treatment in aged mice (Fig. 5E).

**Figure 5:**
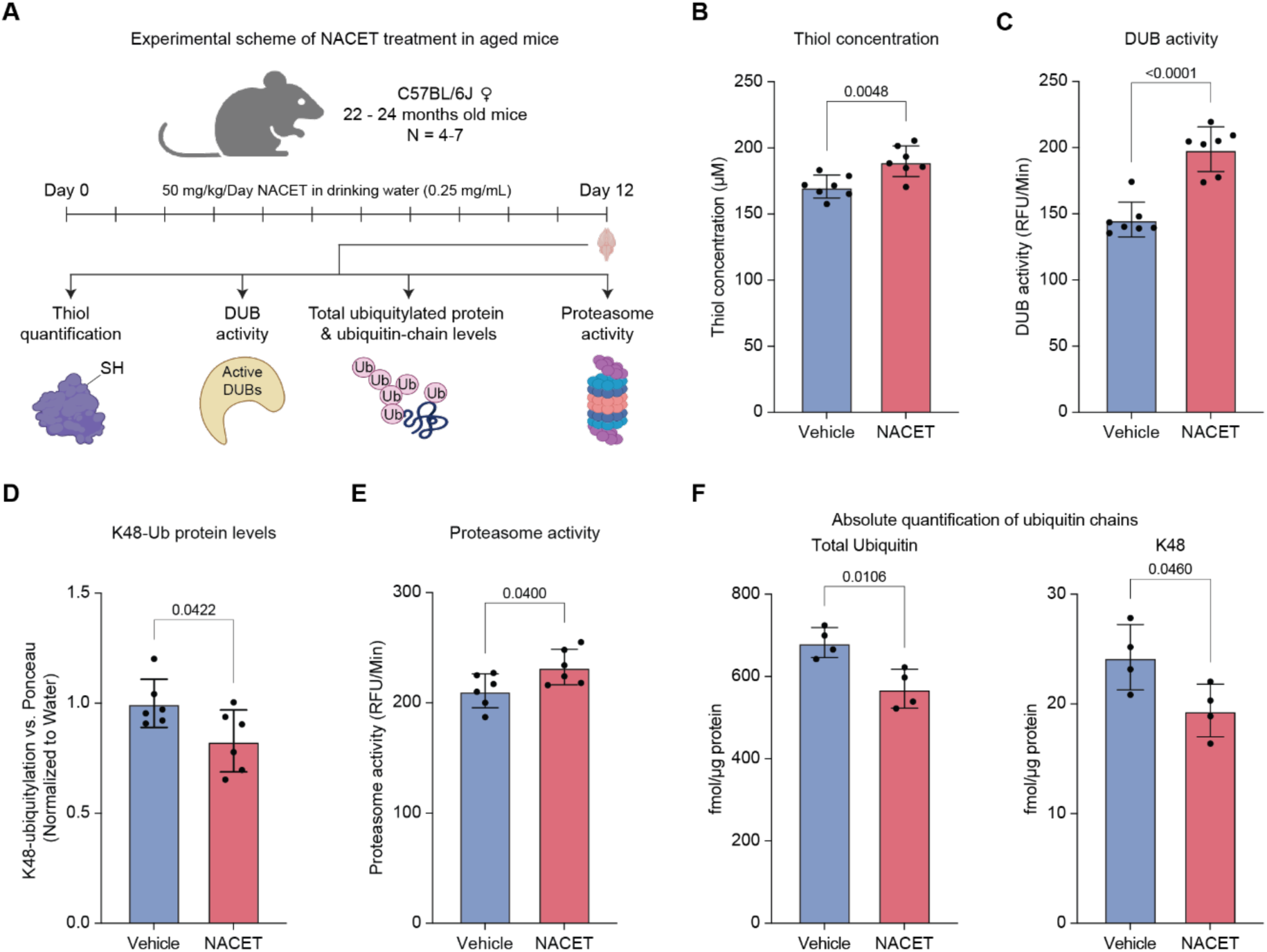
NACET reverses age-associated redox and proteostasis alterations. (A) Schematics of N-acetylcysteine ethyl ester (NACET) treatment in C57BL/6J aged 22 - 24-month-old female mice. Animals were treated with 0.25 mg/mL NACET in drinking water to achieve a final concentration of 50 mg/kg/day of NACET (N = 4-7 biological replicates, mice used from two independent experimental cohorts). (B) Reduced thiol concentrations in the NACET-treated mouse brains (N = 7). (C) DUB activity (N = 7). (D) K48 polyubiquitylated protein abundance quantification normalized to total protein levels (N = 6). (E) Proteasome activity (N = 6). Data in all panels are pooled from two independent experiments, with technical outliers excluded as described in the Methods section. (F) Absolute quantification (AQUA-PRM) of total ubiquitin and K48-linked ubiquitin chains in NACET vs. vehicle-treated aged mouse brains (N = 4 biological replicates). In all panels, an unpaired t-test with Welch’s correction was used. RFU represents Relative Fluorescence Units. Related to Supplementary Tables 1 and 4.

**Supplementary Figure 5:**
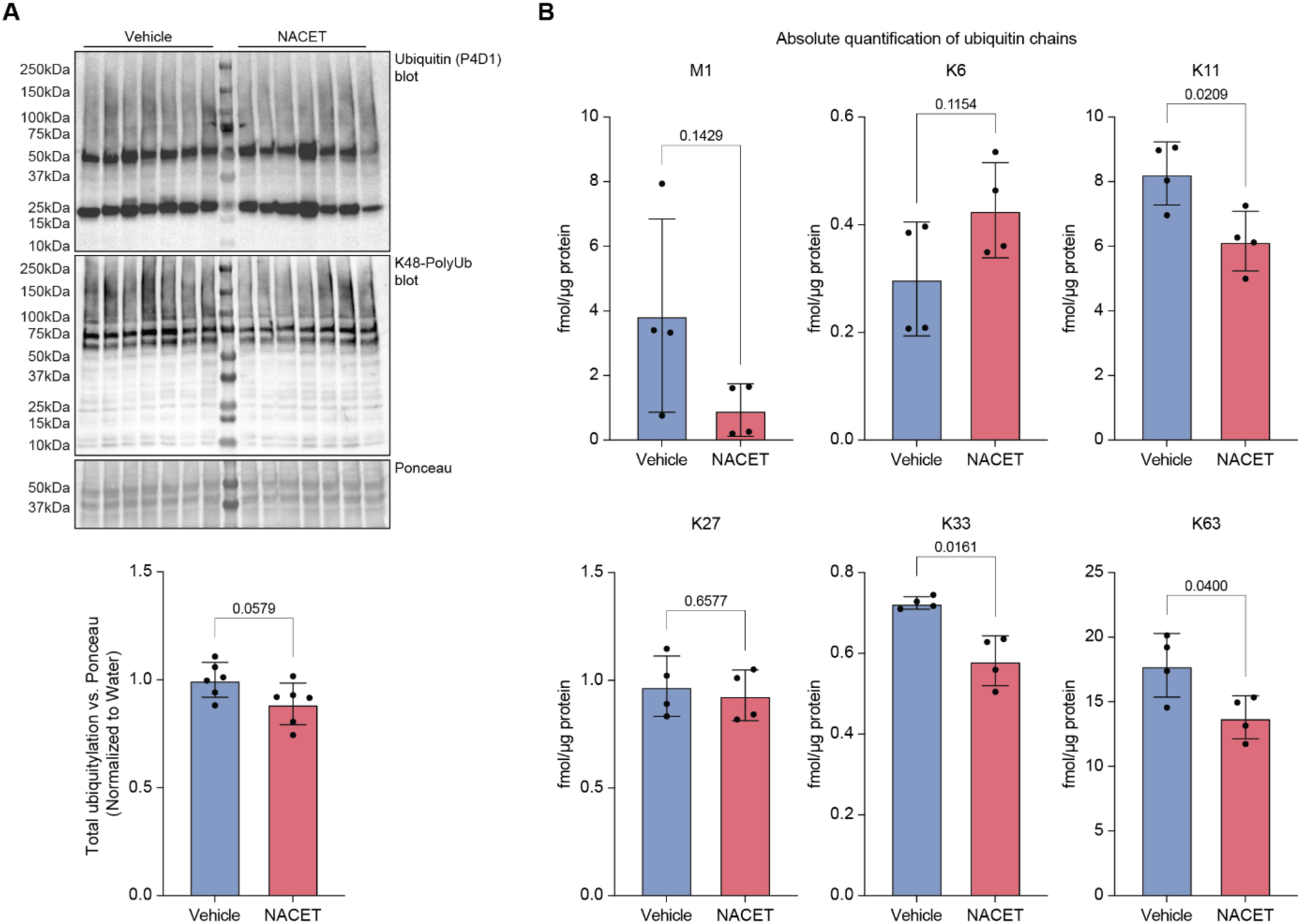
NACET treatment rescues brain ubiquitylation in aged mice. (A) Upper: Immunoblot of total and K48 polyubiquitylated protein levels. Total protein levels were assessed using Ponceau staining. Lower: Total ubiquitylated protein abundance quantification normalized to total protein levels. Replicate 1 from vehicle and replicate 4 from NACET-treated animals were removed from both quantification analyses (N = 6 biological replicates). (B) AQUA-PRM of ubiquitin chain linkages in NACET vs. vehicle-treated aged mouse brains (N = 4 biological replicates; unpaired t-test with Welch’s correction). Related to Supplementary Table 4.

Since aging alters specific ubiquitin-chain linkage abundances in mouse brain ^5^, we next examined whether NACET could restore these molecular changes beyond proteostasis. Using targeted proteomics via parallel reaction monitoring with absolutely quantified isotopically labeled spike-in reference peptides (AQUA-PRM) (Fig. 5A), we found that NACET treatment decreased total ubiquitin and K48 chain levels (Fig. 5F), consistent with immunoblot results. Additionally, NACET reduced K11, K33, and K63 chain abundances, which are known to increase during brain aging (Fig. S5B). Taken together, these findings demonstrate that NACET treatment restores redox homeostasis in aged brains, leading to reactivation of DUB activity, reduction of deleterious polyubiquitin chain accumulation, and enhancement of proteasome function, thereby improving overall protein quality control during brain aging.

## Discussion

Aging is associated with a gradual decline in protein homeostasis, which contributes to the onset and progression of neurodegenerative diseases. While proteasomal dysfunction has been extensively studied in the context of aging and age-associated pathologies ^2,3,48–52^, upstream regulators of ubiquitin turnover, such as deubiquitylating enzymes (DUBs), remain less explored. In this study, we systematically profiled the activity landscape of cysteine protease DUBs during brain aging in both mouse and killifish, uncovering a redox-sensitive subset of enzymes that lose almost 40% of their catalytic activity with age despite unchanged protein abundance. Notably, several DUBs identified in our study - including ATXN3, UCHL1, UCHL5, and YOD1 - have been previously implicated in neurodegenerative disorders such as spinocerebellar ataxia, Alzheimer’s disease, and Parkinson’s disease ^10,20–22,53,54^, underscoring the relevance of DUB activity loss in the context of aging and age-related neurodegeneration.

Mitochondrial dysfunction and impaired proteostasis - two hallmarks of aging and neurodegeneration - are tightly interconnected ^1,27,55^. For example, mitochondria depend on proper protein import, folding, and degradation to function efficiently; failure in these processes can trigger mitochondrial dysfunction. In turn, dysfunctional mitochondria produce elevated levels of reactive oxygen species (ROS), which can oxidize amino acid residues such as cysteine and methionine. This leads to protein misfolding and impairs key enzymes involved in maintaining proteostasis, ultimately resulting in protein aggregation ^56–59^.

Our findings suggest that such ROS-driven thiol oxidation underlies the age-dependent loss of DUB activity in aged mouse brains, which was reversible upon *in vitro* treatment with dithiothreitol (DTT), a reducing agent. Furthermore, *in vivo* administration of the antioxidant N-Acetyl-L-cysteine ethyl ester (NACET) increased the reduced thiol pool and enhanced DUB activity in old animals. Our data are supported by the findings of ^11^ and ^60^, which suggest reversible inactivation of cysteine protease DUBs like USP1 by ROS. These results implicate thiol oxidation as a key mechanism suppressing DUB function in the aging brain, disrupting the fine-tuned regulation of protein ubiquitylation, and contributing to proteostasis decline. Although thiol oxidation emerges as a central regulatory mechanism, other modes of DUB modulation - such as phosphorylation, ubiquitylation, and protein-protein interactions - remain to be explored in the context of aging, as comprehensively reviewed by ^6^.

Functionally, we demonstrate that inhibition of DUBs in human iPSC-derived neurons (iNeurons) recapitulates some of the ubiquitylation patterns observed in aged brains ^5^, establishing a causal link between DUB inactivation and age-related proteome changes. For example, alcohol dehydrogenase 5 (ADH5) - one of the proteins showing the most substantial increase in ubiquitylation during aging and following DUB inhibition - is critical for formaldehyde detoxification via a glutathione-dependent mechanism ^61^. Impaired ADH5 function and formaldehyde toxicity have been linked to mitophagy, aging, and age-related neurological disorders ^40,61,62^, suggesting that its post-translational modification may affect protein stability or function.

Likewise, proteins involved in cytoskeletal dynamics, proteolysis, and protein folding exhibited increased ubiquitylation following DUB inhibition and in aged mouse brains, as previously described by us ^5^. Notably, Heat Shock 70 kDa Protein 8 (HSPA8) - a chaperone whose reduced protein levels have been associated with the accumulation of neurodegeneration-related proteins and increased oxidative stress ^63,64^ - shows increased ubiquitylation at most lysine residues during aging. However, specific sites such as K56 and K126 display reduced ubiquitylation in both aged brains ^5^ and Huntington’s disease models ^65^. Both K56 and K126 reside in the ATPase domain of HSPA8, and their post-translational modification has been implicated in modulating, particularly inhibiting chaperone activity ^66–68^. The fact that these site-specific changes are reproduced by DUB inhibition but not by proteasome inhibition ^5^, highlights the distinct role of DUBs in maintaining HSPA8 chaperone activity by regulating its ubiquitylation.

Conversely, we also observed decreased ubiquitylation of a subset of proteins involved in synaptic function and mRNA processing upon DUB inhibition in iNeurons. Proteins (such as Epsin 1 and CAMK2A, see Supplementary Table 3) belonging to the same functional categories show reduced ubiquitylation levels during aging ^5^ and depolarized synaptosomes ^69,70^. One possible explanation for such cases is an indirect regulatory effect, where disrupted DUB activity alters the coordinated interplay between DUBs and E3 ligases. For instance, inhibition of USP2a or USP7 has been shown to reduce p53 ubiquitylation by destabilizing the E3 ligase MDM2 ^71,72^. This highlights the importance of DUB-E3 interactions in shaping the ubiquitylated proteome landscape. A comprehensive analysis of how (E1, E2, and) E3 ligase activity is modulated during aging and how this contributes to age-associated proteome remodeling and disease remains an important future direction. Tools such as Ub-Dha-based activity probes could be instrumental in mapping these dynamics ^73^.

Building on our findings of decreased DUB activity (this study) and proteasome decline during aging ^3,5^, we next sought to determine the sequence of molecular events leading to proteostasis disruption in the aging brain. A temporal analysis across different mouse age groups revealed a progressive, thiol oxidation-mediated reduction in DUB activity from middle to old age (12-18 months). In contrast, proteasome activity varied with age and showed a significant decline only in later life, consistent with observations of 20S proteasome fluctuations during aging reported by ^74^. Furthermore, in line with findings from ^75^ that reported reduced proteasome activity in SH-SY5Y dopaminergic cells following USP14 knockout, and ^45^, that reported impaired proteasome function in OLN-t40 oligodendroglial cells upon global DUB inhibition by PR619, we observed diminished chymotrypsin-like proteasome activity following chronic (24 hours) DUB inhibition in iNeurons. Collectively, these results suggest that DUB dysfunction is an early molecular event in the proteostasis cascade, preceding and potentially contributing to proteasome impairment during aging.

Finally, we have demonstrated that NACET treatment not only restored DUB activity but also rescued proteasome function in aged mouse brains. This effect may be explained by three potential mechanisms: (1) enhanced DUB activity reverses stress-induced ubiquitylation on proteasome subunits such as RPN10 or RPN13, which otherwise promotes dissociation of the 19S regulatory particle from the 26S complex. This autoinhibitory mechanism is thought to have evolved as a protective strategy to prevent the engagement of ubiquitylated substrates with defective or stalled proteasomes, thereby minimizing futile degradation attempts under stress conditions ^76–78^; (2) by relieving oxidative stress, NACET may independently promote 26S complex reassembly and enhance ATPase activity, thereby restoring proteasome function ^79–82^; or (3) both mechanisms may act in parallel to achieve the observed rescue effect. These findings highlight the potential of redox interventions in late life to restore aspects of proteostasis and underscore the reversibility of oxidative inactivation. However, the long-term physiological consequences and therapeutic implications of such interventions remain to be systematically explored in both aging and disease-relevant models.

Although we report an age-dependent decline in DUB activity that may negatively impact proteostasis components and metabolic enzymes, other studies suggest that DUB inhibition can have beneficial effects in the context of aging and disease. For example, ^4^ observed reduced levels of ubiquitylated proteins during aging in *C. elegans*, which could be rescued by global DUB inhibition using the broad-spectrum inhibitor PR619 ^32^. This apparent discrepancy may reflect species-specific (vertebrate vs. invertebrate) or tissue-specific (brain vs. whole organism) or cell type-specific differences ^5,83–85^. Notably, *C. elegans* encodes only about half as many DUBs as vertebrates. While some DUBs have conserved functions across species, many differ in substrate specificity, regulatory pathways, and mechanisms of action, potentially explaining the divergent outcomes of DUB modulation.

Moreover, several studies have demonstrated that genetic or pharmacological inhibition of proteasome-associated DUBs, such as USP14 and UCHL5, can enhance proteasome activity across various systems, including *in vitro* assays, cultured cells, and *C. elegans* models *^42,43,86^*. Similarly, inhibition of USP30 has demonstrated neuroprotective effects in Parkinson’s disease models ^87,88^, while USP7 inhibition has been associated with reduced neuroinflammation ^89^. Numerous other DUBs have also been implicated in neurodegenerative contexts ^90,91^, as comprehensively reviewed by ^92^.

Importantly, the reported increase in proteasome activity and enhanced clearance of toxic proteins upon DUB inhibition appears to be highly context-dependent. These effects are most consistently observed in *C. elegans*, cancer cell lines, or disease models characterized by abnormal protein accumulation. Such benefits may be mediated through mechanisms including 20S proteasome gate opening, reduced ubiquitin chain trimming, or non-catalytic regulatory effects of DUBs on proteasome function ^44^.

In conclusion, we identify a subset of DUBs that undergo selective and reversible inactivation with age due to thiol oxidation. This loss of deubiquitylation capacity emerges as an early molecular feature of aging that precedes proteasomal decline and contributes to proteostasis disruption. Our findings provide mechanistic insight into the hierarchical dysfunction of the ubiquitin system in the aging brain and suggest novel opportunities to restore proteostasis by preserving cysteine proteases DUB function.

## Methods

### Mice

Wild-type mice (*Mus musculus*) were either C57BL/6J or C57BL/6N, obtained from Janvier Labs or internal breeding at FLI. Animals were kept in a specific pathogen-free animal facility with unlimited access to food and water at FLI, Jena (C57BL/6J) or ZMG, Halle (C57BL/6N). The housing conditions were adjusted to a 12-hour day-night cycle, temperature and humidity of 20 °C ± 2 and 55% ± 15, respectively. The following aged male mice were used in the experiment and are mentioned in each figure: 3, 6, 12, 18, 24, and 30 months old. Female mice of 22 - 24 months were used. In all figures, N refers to the number of biological replicates used. C57BL/6J mice were euthanized with CO_2,_ whereas C57BL/6N mice were sacrificed by cervical dislocation (approved code: K2bM3). Brains were isolated, washed in PBS, cut into two halves, weighed, immediately snap-frozen in liquid nitrogen or dry ice, and stored at -80 °C. The guidelines from the European Parliament on the protection of animals, Directive 2010/63/EU, and the guide for the care and use of laboratory animals, 8th edition, 2011, Washington (DC), were used for all experiments. Sacrifice and organ collection were carried out following §4(3) of the German Animal Welfare Act.

### Killifish

All experiments were conducted using the male *Nothobranchius furzeri* strain MZM-0410 (N = 3 biological replicates) in accordance with institutional and national guidelines. Fish were bred and maintained in the FLI fish facility in accordance with §11 of the German Animal Welfare Act. Sacrifice and organ collection were carried out following §4(3) of the German Animal Welfare Act. Fish were housed individually or in groups (maximum one fish per 1.7 L) in recirculating systems (Aqua Schwarz, Göttingen, Germany) with centralized filtration, maintained under a 12-hour day-night cycle. Water temperature (27 ± 1 °C), conductivity (∼2.5 mS), and pH were continuously monitored and recorded. As environmental enrichment, group-housed fish were provided with certified contaminant-screened blue polycarbonate igloos and shelters (Bio-Serv, Flemington). Newly hatched larvae were fed twice daily with live *Artemia nauplii* and weaned onto live bloodworms (*Chironomidae*) once daily from 4 to 6 weeks of age. Fish health was assessed daily by trained caretakers using the FLI killifish health score sheet ^93^, and animals were euthanized by rapid chilling if humane endpoints were reached. Health monitoring followed the protocols described in ^93^.

### NACET treatment in aged mice

For N-acetylcysteine ethyl ester (NACET) treatments on aged mice, 22 to 24-month-old female C57BL/6J mice were used. Following established protocols ^46,47^, NACET was resuspended in drinking water at 0.25 mg/mL. Then, based on the average daily water intake of ∼5 mL per mouse, 50 mg/kg of NACET was administered daily to each animal, and a freshly prepared NACET solution was replaced every 2 days. Control mice received regular drinking water. After 12 days of treatment, mice were sacrificed, and brains were collected as described above and stored at - 80 °C for downstream analyses. The experiment was performed independently twice, with N = 3 and N = 4 biological replicates per experiment. Animal experiments were approved by the Thuringian State Office for Consumer Protection (Thüringer Landesamt für Verbraucherschutz, license number: FLI-24-011).

### Activity-based DUB probe labeling in mouse and killifish brains

Mouse and killifish brains were lysed in DUB buffer consisting of 50 mM Tris-HCl (pH 7.5), 150 mM NaCl, 0.1% NP-40, 5% CHAPS, 5 mM MgCl₂, 5 mM β-mercaptoethanol, and 10% glycerol. Killifish brains were lysed using a vial tweeter (Amplitude 100%, cycle 0.9), while mouse brains were manually disrupted by mechanical trituration. This was performed by repeatedly aspirating and dispensing the tissue using a 2 mL syringe (Injekt) fitted first with a 0.70 × 30 mm needle, followed by a 0.45 × 25 mm needle (Sterican). After lysis, samples were subjected to a short centrifugation step to pellet debris, and the supernatants were transferred to fresh 1.5 mL centrifuge tubes. Protein concentrations were determined by measuring absorbance at 280 nm using a NanoDrop 2000 spectrophotometer (Thermo Scientific). Mouse brain lysates equivalent to 230 µg of protein were either reduced with 10 mM dithiothreitol (DTT) or mock-treated with DUB buffer (control) for 30 minutes at 300 rpm and 25 °C. Both mouse and killifish lysates were subsequently alkylated with 10 mM N-ethylmaleimide (NEM; Thermo, Cat# 23030) for 30 minutes at 300 rpm and 25 °C to block free sulfhydryl groups, serving as a negative control for activity-based probe labeling.

Following NEM or DMSO treatment, samples were incubated with a 2.3 µM mixture of three activity-based probes: Biotin-Ahx-Ub-VME (UbiQ-054), Biotin-Ahx-Ub-PA (UbiQ-076), and Biotin-Ahx-Ub-VS (UbiQ-188), for 60 minutes at 300 rpm and 25 °C. The reaction was terminated by adding 0.4% SDS, followed by freezing and storage at -20 °C. The volume corresponding to ∼30 µg of protein was taken out from each reaction. Of this, 10 µg was used for immunoblotting to validate probe labeling, and 20 µg was stored at -20 °C for future use. Excess unbound probe was removed from the remaining reaction mixture (∼200 µg protein) using 10 kDa Amicon Ultra centrifugal filters (Merck, UFC5010) with DUB buffer containing 2% SDS.

### Enrichment and Digestion of Activity-Labeled DUBs

To enrich DUBs labeled by the activity-based probes, an automated workflow using the Agilent Bravo AssayMAP platform was employed, with minor modifications based on ^94^. Streptavidin cartridges were first equilibrated with 200 µL of PBS at 10 µL/min and acetylated with 50 µL of 10 mM sulfo-NHS acetate. Before sample loading, the cartridges were equilibrated with 300 µL of DUB buffer (internal cartridge wash 1) at 20 µL/min. Filtered lysates were loaded at 10 µL/min. After binding, the cartridges were washed once with 250 µL of DUB buffer containing 2% SDS and twice with 250 µL of 50 mM ammonium bicarbonate (AmBic) at 10 µL/min. Proteins were digested on-cartridge using 0.5 µg of LysC (Cell Signaling) in 30 µL of 50 mM AmBic. Digestion was performed in a 6 µL reaction volume at 45 °C for 60 minutes without a reaction chase. Peptides were eluted in two steps using 100 µL of 50 mM AmBic (no internal cup wash, 10 µL/min). Eluates were subsequently digested with trypsin (0.5 µg/µL, Promega) at 37 °C and 500 rpm overnight. Reactions were acidified with 10% trifluoroacetic acid (TFA) to a pH < 3. Peptides were desalted using Waters Oasis® HLB µElution Plate (30 µM) according to the manufacturer’s instructions and sent for DUB identification and quantification using LC-MS/MS, as mentioned below.

### Mouse brain homogenization

Snap-frozen half-brains stored at -80 °C were thawed on ice and transferred to a 5 mL Douncer. The volume of pre-chilled PBS added to each brain sample was calculated based on the estimated protein content (∼5% of the fresh tissue weight) to reach a 20 µg/µL concentration. Homogenization was performed for 15-20 passes of the pestles up and down the glass cylindrical Douncer, followed by 15 seconds of spin. The homogenization step was repeated for three cycles, and the total homogenate was transferred to a 1.5 mL centrifuge tube. The homogenates were clarified from tissue debris by centrifuging at 17,000 × g for 1 minute at 4 °C, and the clarified homogenate was aliquoted to multiple 1.5 mL centrifuge tubes. The aliquots were then stored at -80 °C for future experiments.

### DUB activity assay

To 5 µL of brain homogenates, 75 µL 1× DUB assay buffer (without DTT), supplied with Abcam’s Deubiquitinase Assay Kit (ab241002), was added. The lysates were prepared by sonication for 60 sec ON/ 30 sec OFF at 4°C for five cycles in a Bioruptor Plus sonicator at high intensity setting. The lysates were centrifuged at 10,000 × g for 5 minutes at 4°C, and the supernatants were then transferred to a new 1.5 mL centrifuge tube. The protein concentration was measured at A280 using Thermo Scientific NanoDrop 2000. Lysates corresponding to 5 µg of protein were used to measure DUB activity as per the manufacturer’s protocol. As a negative control, NEM was used at a final concentration of 10 mM per reaction. Fluorescence was measured after setting up the reaction in the kinetic mode for 60 minutes at 25°C by TECAN kinetic analysis (excitation 350 nM, emission 440 nM, 30-second reading interval) on a Safire II microplate reader (TECAN). The DUB activity was determined as the difference between the activity of protein lysates and the residual activity of the lysate in the presence of NEM, and results were represented as relative fluorescence units per minute (RFU/Min). Similar steps were followed for the iNeurons lysate treated with DMSO, PR619, or Bortezomib, except that RFU was calculated from the difference between the activity of protein lysates in the presence and absence of DUB substrates. Outliers were removed based on raw fluorescence data, if pipetting errors or air bubbles were identified after measurement, or if the final RFU value fell outside one standard deviation from the mean.

### Proteasome chymotrypsin-like activity assay

To 7 µL brain homogenates, 50 µL 1× lysis buffer consisting of a final concentration of 1× Proteasome assay buffer (supplied with the UBPBio’s Proteasome Activity Fluorometric Assay Kit II, J4120), 50 mM NaCl, 2 mM ATP, 5 mM MgCl2, and 10% Glycerol. While for the iNeurons pellet, 100 μL 1× lysis buffer was used. The lysates were prepared, and protein concentration was measured as mentioned in the DUB activity assay methodology. The final concentration of 1× Suc-LLVY-AMC fluorogenic peptides supplied with the kit was used to measure the chymotrypsin-like activity (CT-L) of the proteasome using 50 µg of proteins in a total of 100 µL reaction as per the manufacturer’s protocol using a 96-well plate (Falcon, 353219). Fluorescence was measured after setting up the reaction in the kinetic mode for 60 minutes at 37°C by TECAN kinetic analysis (excitation 360 nM, emission 460 nM, 30-second reading interval) on a Safire II microplate reader (TECAN). The CT-L activity was determined as the difference between the activity of protein lysates and the residual activity of the lysate in the presence of 100 µM MG132 supplied with the kit, and results were represented as Relative Fluorescence Units per minute (RFU/Min). Similar steps were followed for the iNeurons lysate treated with DMSO, PR619, or Bortezomib, except that RFU was calculated from the difference between the activity of protein lysates in the presence and absence of fluorogenic peptides. Outliers were removed based on raw fluorescence data, if pipetting errors or air bubbles were identified after measurement, or if the final RFU value fell outside one standard deviation from the mean.

### Thiol concentration measurement using DTNB assay

To 15 µL of brain homogenate, 50 µL of reaction buffer was added. The reaction buffer contained 0.1 M sodium phosphate dibasic (Sigma, S0876) and 1 mM EDTA (Roth, 8040.3) at pH 8.0. The lysate preparation and protein concentration measurement were conducted as outlined in the DUB activity assay methodology. For the experiment, lysates corresponding to 100 µg of proteins were combined with 5 µL of 5,5’-dithio-bis-[2-nitrobenzoic acid] (DTNB, Thermo Scientific, 22582) to achieve a total volume of 280 µL in a 96-well plate (Falcon, 353219). L-Cysteine (Sigma, C7352) standards were prepared at concentrations of 0, 10, 50, 100, 200, 400, 800, and 1000 µM in the reaction buffer, and set up the reaction as experimental samples using DTNB. The reactions were then incubated for 15 minutes at 22°C. The absorbance was measured at 412 nm using the Safire II microplate reader (TECAN). The experimental samples were corrected using non-treated lysates as blanks for measuring thiol concentration. Based on the absorbance vs. cysteine concentration trendline curve equation derived from the L-cysteine standards, the thiol concentration for the unknown experimental samples was extrapolated. Outliers were removed based on raw fluorescence data, if pipetting errors or air bubbles were identified after measurement, or if the final thiol concentration values fell outside one standard deviation from the mean.

### Immunoblot analysis

Samples for immunoblot analysis were used as prepared and mentioned previously. Otherwise, mouse brain homogenate or iNeurons pellets were lysed in 1× lysis buffer containing 2% SDS and 25 mM HEPES for 60 sec ON/ 30 sec OFF at 20°C for ten cycles in a Bioruptor Plus sonicator at high-intensity setting. Lysates were boiled at 95°C for 5 minutes, followed by centrifugation for 1 minute at 17,000 × g. Supernatants were transferred to new 1.5 mL centrifuge tubes, and protein concentrations were measured using Pierce BCA Protein Assay Kit (Thermo Scientific, 23225). Subsequently, samples were reduced using DTT (Carl Roth, 6908) at a final concentration of 10 mM for 15 minutes at 45 °C. 10-20 µg of proteins were used along with 4x sample loading buffer containing 1.5 M Tris pH 6.8, 20% (w/v) SDS, 85% (v/v) glycerin, 5% (v/v) β-mercaptoethanol. The sample mixture was then incubated at 95 °C for 5 minutes. For immunoblots, proteins were separated on 4-20% Mini-Protean® TGX™ Gels (Bio-Rad #4561096) by sodium dodecyl sulfate-polyacrylamide gel electrophoresis (SDS-PAGE) using a Mini-Protean® Tetra Cell system (Bio-Rad, Neuberg, Germany, 1658005EDU). Proteins were then transferred using a Trans-Blot® Turbo™ Transfer Starter System (Bio-Rad #1704150) onto nitrocellulose membrane (Carl Roth, 200H.1). For measurement of total protein levels on blot, membranes were stained with Ponceau S (Sigma, P7170-1L) for 5 minutes on a shaker (Heidolph Duomax 1030), followed by washing with Milli-Q water. The Ponceau staining was then imaged on a Molecular Imager ChemiDocTM XRS + Imaging system (Bio-Rad) and destained by three washes in TBST (Tris-buffered saline (TBS, 25 mM Tris, 75 mM NaCl), with 0.5% (v/v) Tween-20) for 10 minutes each. The blots were then incubated in PBS for 1 hour in a house-made blocking buffer of 3% BSA (w/v) and 0.5% Tween 20 (v/v). Post-blocking blots were incubated overnight at 4°C on a tube roller (BioCote® Stuart® SRT6) with primary antibodies against total ubiquitin P4D1 (1:1000, Santa Cruz #sc8017), Lys-48 specific anti-ubiquitin antibody (1:1000, Sigma Aldrich #05-1307), or Streptavidin-HRP (1:20,000, Abcam #ab7403) in an enzyme dilution buffer containing 0.2% BSA (w/v) and 0.1% Tween20 (v/v) in PBS. The next day, blots were washed three times with TBST for 10 minutes each at room temperature, and Streptavidin-HRP blots were imaged while ubiquitin blots were incubated with horseradish peroxidase-coupled secondary antibody (Dako #P0447 or #P0448) at room temperature for 1 hour (1:2000 in 0.3% (w/v) BSA in TBST). Following incubation with secondary antibody, blots were washed 3 times for 10 minutes in TBST. The chemiluminescent signals were detected using an ECL Pierce detection kit (Thermo Fisher Scientific, Waltham, MA, USA, 32109) on the Molecular Imager ChemiDocTM XRS + Imaging system (Bio-Rad). The results were analyzed using Bio-Rad’s Image Lab 6.1 software. For protein normalization, Ponceau was used.

### Human iPSC maintenance

The WTC11 human induced pluripotent stem cells (iPSCs) were maintained according to ^33^ and were a kind gift from the Ward Lab at the National Institute of Health, US. Briefly, the tissue culture surface was coated with 1× Matrigel (Corning, 356231) coating solution prepared in DMEM/F12 (Gibco, 11320033) medium for at least 1 hour and then removed from the surface. iPSCs were thawed from liquid nitrogen and washed once with DMEM/F12, followed by resuspended in Essential 8 (E8) culture medium (Gibco, A1517001) supplemented with Y-27632 ROCK inhibitor (Abcam, ab120129) (Day 1). The cells were incubated at 37 °C. The culture was maintained by daily E8 medium change without the presence of 10 µM ROCK inhibitor. Every fourth day (Day 4), cells were split by detaching using Accutase (Sigma, A6964), resuspended in E8 medium supplemented with ROCK inhibitor, and seeded into a new Matrigel-coated surface. For storage in liquid nitrogen, Day 4 cells were frozen in cryopreservation medium containing 20% FBS and 10% DMSO in E8 medium.

### iPSC differentiation to iNeurons and drug treatment regimen

Once iPSCs reached 70%-80% confluency (Day 4 of iPSC), they were detached using Accutase, pelleted down at 300 × g for 5 minutes. The cell pellets were then resuspended and transferred to a Matrigel-coated surface in induction medium (IM) (DMEM/F12 (Gibco, 11320033) supplemented with N2 supplement (Gibco, 17502048), L-glutamine (Gibco, 25030081), MEM NEAA (Gibco, 11140050), and 2 µg/mL doxycycline (Sigma, D9891)) supplemented with 10 µM Y-27632 ROCK inhibitor (Abcam, ab120129). The cells were incubated at 37 °C (Day 1 of iNeurons). The next day, the culture was maintained by changing the IM medium without ROCK inhibitor (Day 2). On the same day, new plates were coated with 1 mg/mL Poly-L-ornithine hydrobromide (Sigma Aldrich, 27278-49-0) in the buffer containing 100 mM boric acid, 75 mM sodium chloride, 25 mM sodium tetraborate, and 1 M sodium hydroxide (Day 2). A day after (Day 3), the PLO-coating was removed from the surface, washed three times with PBS, and dried under the hood for at least 30 minutes. Meanwhile, differentiated cells were dissociated from the Matrigel-coated plate’s surface using Accutase, washed with DMEM/F12, and pelleted down. The cells were then resuspended in cortical neuron culture medium (CM) consisting of Neurobasal medium (Gibco, 10888022) supplemented with B-27 supplement (Gibco, 17504044), 10 ng/mL NT-3 (PeproTech, 450-03), 10 ng/mL BDNF (PeproTech, 450-02), 1 µg/mL laminin (Sigma-Aldrich, L2020), 2 µg/mL doxycycline (Sigma, D9891), and 10 µM Y-27632 ROCK inhibitor and seeded to the PLO-coated surface. The iNeurons were then cultured by removing half of the old medium and exchanging it with fresh CM medium without ROCK inhibitor biweekly. For long-term storage in liquid nitrogen, Day 3 differentiated iNeurons were resuspended in 20% FBS and 10% DMSO in CM media and aliquoted as 6 million/ mL per vial.

The DUB and proteasome inhibitions experiments were performed on 14-day post-differentiated iNeurons. PR619 (Sigma, 662141) was used to inhibit cysteine DUBs, while the proteasome was inhibited using Bortezomib (Sigma, 5043140001). Drugs were prepared in DMSO and treated to iNeurons by removing half of the old medium and exchanging it with fresh CM medium with double the required final drug concentrations. In all experiments, the final concentration of Bortezomib was 10 nM, and the cells were treated for 24 hours. 10 µM PR619 for 6 hours was used to inhibit DUBs in the ubiquitylated peptides enrichment experiment. 5, 7.5, and 10 µM PR619 for 24 hours were used to assess the impact of DUB inhibition on the proteasome. For cell harvesting, iNeurons were washed three times with chilled PBS. Accutase was added to the cells and incubated for 10 minutes at 37 °C. After 10 minutes, the reaction was stopped by adding DMEM/F12. Using a cell scraper, iNeurons were scraped gently from the surface, collected into 1.5 mL centrifuge tubes, and pelleted down at 1,000 × g for 1 minute. Pellets were then stored at -80°C for downstream experiments.

### Sample preparation for total proteome and analysis of ubiquitylated peptides

Brain homogenates corresponding to 1.1 mg and iNeurons pellets were lysed in 1× lysis buffer containing 2% SDS and 25 mM HEPES final concentrations for 60 sec ON/ 30 sec OFF at 20°C for ten cycles in a Bioruptor Plus sonicator at high-intensity setting. Lysates were boiled at 95°C for 5 minutes, followed by centrifugation for 1 minute at 17,000 × g. Supernatants were transferred to new 1.5 mL centrifuge tubes, and protein concentrations were measured using Pierce BCA Protein Assay Kit (Thermo Scientific, 23225). Subsequently, samples were reduced using DTT (Carl Roth, 6908) at a final concentration of 10 mM for 15 minutes at 45 °C. Proteins corresponding to 1.1 mg were then alkylated using freshly prepared iodoacetamide (IAA) (Sigma-Aldrich, I1149) with a final concentration of 15 mM for 30 minutes at room temperature in the dark. Cold acetone corresponding to 4 times the volume was added to samples and incubated overnight at -20°C, as described in ^95^. The next day, acetone was removed post-centrifugation at 17,000 × g at 4°C, washed twice with 80% ice-cold acetone, and resuspended in digestion buffer consisting of 3 M urea and 100 mM HEPES at pH 8.0 such that the final protein concentration was 1 µg/µL. Subsequently, proteins were digested using 1:100 (enzyme: protein) LysC (Wako sequencing grade, 125-05061) at 37 °C for 4 hours, followed by dilution with HPLC water to make a 1.5 M final urea concentration and digestion with 1:100 Trypsin (Promega sequencing grade, V5111) for 16 hours. The next day, the reaction was stopped by acidifying peptides with 10% trifluoroacetic acid (v/v). Peptides corresponding to 20 µg and 1000 µg were taken for proteome and ubiquitylated peptides enrichment, respectively. Peptides were desalted using Waters Oasis® HLB µElution Plate (30 µM, 2 mg for proteome and 30 mg for ubiquitylome) following the manufacturer’s instructions. The cleaned peptides were then dried using a vacuum concentrator at 45 °C and reconstituted in 5% acetonitrile (v/v) and 0.1% formic acid (v/v) MS Buffer A. For total proteome analysis, 20 µg of diluted peptides at 1 µg/µL were transferred to MS vials, spiked with iRT kit peptides (Biognosys, Ki-3002-2), and sent for LC-MS/MS. Meanwhile, samples were further processed for ubiquitylated peptide enrichment, as described below.

### Enrichment of ubiquitylated peptides

Dried peptides corresponding to ∼1000 µg were used to enrich ubiquitylated peptides. The PTMScan® HS Ubiquitin/SUMO Remnant Motif (K-ε-GG) kit (Cell Signaling Technology, 59322) was used, and peptides were enriched according to the manufacturer’s instructions. Thereafter, the enriched K-ε-GG modified peptides were desalted, concentrated, and prepared in MS vials for the LC-MS/MS analysis as described above.

### Absolute quantification of ubiquitin chain linkages

Followed the standardized protocol for parallel reaction monitoring (PRM)-based measurement of endogenous ubiquitin chain linkages using Absolute QUAntification (AQUA) synthetic peptides, as described by us before ^5^, with minor modifications. Briefly, following acetone precipitation and resuspension in digestion buffer, protein samples corresponding to 20 µg in 20 µL were obtained from NACET- and vehicle-treated mouse brain lysates. AQUA peptides were spiked into each sample at a concentration of 20 fmol per 1 µg of estimated protein content prior to digestion. Subsequent processing was performed as described in the “Sample preparation for total proteome and analysis of ubiquitylated peptides” section, with the only modification being the use of 10% formic acid instead of 10% trifluoroacetic acid (TFA) for peptide acidification prior to desalting. Peptides were desalted using a Waters Oasis® HLB µElution Plate (30 µm, 2 mg) and resuspended in 10 µL of MS Buffer A. Peptide concentration was re-assessed using a Thermo Scientific NanoDrop 2000 at A280. The final concentration of endogenous peptides was 0.4 µg/µL (total 4 µg), and AQUA peptide concentration was 40 fmol/µL (total 400 fmol) in 10 µL of MS Buffer A. Peptides were transferred to MS vials, and 3 µL of the peptide mixture was injected into a nanoAcquity UPLC M-Class system coupled to an Orbitrap Fusion Lumos mass spectrometer, following the protocol described in ^5^. Peak group identification was carried out using SpectroDive (12.0.24) and subsequently verified manually. Quantification employed a spike-in strategy, calculating the ratio between endogenous (light) and reference (heavy) peptides to achieve absolute quantification. All AQUA peptides corresponding to total ubiquitin and various linkage types were quantified, with the exception of K29.

### Data-independent acquisition mass spectrometry

For DUB analysis on the Evosep platform and proteome of NACET- and vehicle-treated mouse brains, samples were loaded onto Evotips following the manufacturer’s instructions and using protocols adapted from ^94^. Briefly, peptides were separated using the Evosep One system (Evosep, Odense, Denmark) equipped with either an 8 cm × 150 μm i.d. column packed with 1.5 μm Reprosil-Pur C18 beads (Evosep Performance, EV-1109, PepSep, for mouse DUB profiling and NACET- and vehicle-treated mouse brains proteome) or a 15 cm × 150 μm i.d. column packed with 1.9 μm Reprosil-Pur C18 beads (Evosep Endurance, EV-1106, PepSep, for killifish DUB profiling) for a 44-minute gradient (30 samples per day, 30SPD). Solvent A consisted of water with 0.1% formic acid, and solvent B was acetonitrile with 0.1% formic acid. Liquid chromatography was coupled to an Orbitrap Exploris 480 mass spectrometry (Thermo Fisher Scientific). A detailed description of the mass spectrometry parameters used for data-independent acquisition (DIA) analysis is available in ^94^.

For proteome and ubiquitylome analysis of iNeurons, peptides were separated in trap/elute mode using a nanoAcquity MClass UPLC system (Waters Corporation, Milford, MA, USA) equipped with a trapping column (Waters nanoEase M/Z Symmetry C18, 5 μm, 180 μm × 20 mm) and an analytical column (Waters nanoEase M/Z C18 HSS T3, 1.7 μm, 75 μm × 250 mm). Solvent A consisted of water with 0.1% formic acid, and Solvent B was acetonitrile with 0.1% formic acid. Approximately 1 μL of sample (∼1 μg on-column) was loaded onto the trapping column at 5 μL/min using Solvent A. After a 6-minute trapping step, peptides were eluted through the analytical column at 0.3 μL/min. During the elution step, the gradient of Solvent B increased nonlinearly from 0% to 40% over 120 minutes. The total LC run time was 145 minutes, including column equilibration and conditioning. The LC was interfaced with an Orbitrap Exploris 480 mass spectrometry (Thermo Fisher Scientific, Bremen, Germany) via a Proxeon nanospray source using a Pico-Tip emitter (360 μm OD × 20 μm ID, 10 μm tip; New Objective). The spray voltage was set to 2.2 kV, with the capillary and ion transfer tube temperatures both maintained at 300 °C. The RF ion funnel was set to 30%. For data-independent acquisition (DIA), full MS scans were acquired over an m/z range of 350-1650 at a resolution of 120,000 (FWHM), with a maximum injection time of 60 ms and a 3 × 10⁶ ions AGC target. The default precursor charge state was set to 3^+^. DIA MS/MS scans used 40 variable-width isolation windows across the MS1 range, with stepped normalized collision energies of 25, 27.5, and 30%. MS/MS spectra were acquired at a resolution of 30,000 with a fixed first mass of 200 m/z, using either 35 ms maximum injection time or until 3 × 10⁶ ions were collected. All data were acquired in profile mode using Xcalibur 4.3 and Tune version 2.0 (Thermo).

## Data analysis

### Data processing for mass spectrometry DIA data

Spectral libraries for DUB profiling in mouse and killifish brains, and ubiquitylated peptide and proteome analyses in iNeurons were generated by searching the DIA run using Spectronaut Pulsar (Biognosys, Zurich, Switzerland), as referenced in Table 1. All searches were performed against species-specific protein databases supplemented with a list of common contaminants. Search parameters for variable modifications included Oxidation (M) and Acetyl (protein N-term) across all datasets. In addition, Biotin was included for DUB profiling, while GlyGly (K) was used for ubiquitylome datasets as part of variable modifications. For both ubiquitylome and proteome analysis, carbamidomethylation (C) was used as a fixed modification. Three missed cleavages were allowed for ubiquitylome, whereas two were allowed for the rest of the datasets. Library-based searches were performed with a 1% FDR (false discovery rate) at both the protein and peptide levels. Relative quantification was performed in Spectronaut using the LFQ QUANT 2.0 method with global normalization, precursor filtering percentile utilizing a fraction of 0.2, and global imputation. No imputation was applied for DIA total proteome analysis in iNeurons, and local normalization was used. Relative quantification between conditions was performed using replicate samples and default settings in Spectronaut. Candidate and report tables were exported for downstream analyses.

**Table 1:** MS data used for library generation on Spectronaut Software.

| Dataset | Spectronaut version | Library size |
| --- | --- | --- |
| DUB activity profiling - Killifish | 18.3.23 | 1,047 proteins |
| DUB activity profiling - Mouse | 18.6.23 | 657 proteins |
| Ubiquitylome - iNeurons | 18.7.24 | 19,771 modified peptides (4,786 proteins) |
| Proteome - iNeurons | 18.7.24 | 7,056 proteins |

### Compilation of cysteine DUBs and their interacting partners

Cysteine protease deubiquitylases (DUBs) were compiled using the Mouse Genome Informatics (MGI) database (GO:0101005) from the Jackson Laboratory and the DUBase repository ^17^. Known DUB interactors were annotated based on data from ^16^ and DUBase. When necessary, protein entries were mapped to their human gene name orthologs prior to inclusion or use in downstream analyses.

### GO enrichment analysis

Gene Set Enrichment Analysis (GSEA) was performed using the gseGO function from the clusterProfiler R package ^96^. Protein entries were first mapped to their human gene name orthologues and used as input for the enrichment analysis.

## Acknowledgements

The authors gratefully acknowledge the support of the FLI Proteomics Core Facility, the Mouse and Fish Facilities, and the ZMG Animal Facility. We also thank the Rudolph group at FLI for sharing mice and reagents. We are grateful to Paulius Grigaravicius and Omid Omrani (FLI), as well as Matthias Ebert (ZMG), Doreen Sander, Jennifer Kopietz, and Pascal Rudewig, for their assistance with organ isolation. A.O. is supported by the Else Kröner-Fresenius-Stiftung (award number: 2019_A79), the Fritz Thyssen Foundation (award number: 10.20.1.022MN), the Chan Zuckerberg Initiative Neurodegeneration Challenge Network (award numbers: 2020-221617, 2021-230967, and 2022-250618), and the NCL Stiftung. A.S., T.P., and A.O. are supported by the German Research Foundation (Deutsche Forschungsgemeinschaft, DFG) through the Research Training Group ProMoAge (GRK 2155). F.G. gratefully acknowledges support from the Fondo di Beneficenza Intesa Sanpaolo (project ID: B/2023/0171). The FLI is a member of the Leibniz Association and is jointly funded by the Federal Government of Germany and the State of Thuringia. Some of the figures were created using BioRender. ChatGPT was used for language editing.

## Contributions

Conceptualization: A.K.S., T.P., A.O.

Data curation: A.K.S., A.L.M., D.D.F., A.M.

Investigation: A.K.S., A.L.M., D.D.F., A.M., P.R.W., D.G., C.G.

Methodology: A.K.S., A.L.M., D.D.F

Project administration: T.P., A.O.

Data analysis: A.K.S.

Supervision: F.N., F.G., A.S., T.P., A.O.

Visualization: A.K.S.

Writing-original draft: A.K.S., A.O.

Writing-review & editing: A.L.M., D.D.F., A.M., T.P.

## References

1. López-Otín, C., Blasco, M. A., Partridge, L., Serrano, M. & Kroemer, G. Hallmarks of aging: An expanding universe. Cell 186, 243–278 (2023).

2. Saez, I. & Vilchez, D. The mechanistic links between proteasome activity, aging and age-related diseases. Curr. Genomics 15, 38–51 (2014).

3. Kelmer Sacramento, E., et al. Reduced proteasome activity in the aging brain results in ribosome stoichiometry loss and aggregation. Mol. Syst. Biol. 16, e9596 (2020).

4. Koyuncu, S. et al. Rewiring of the ubiquitinated proteome determines ageing in C. elegans. Nature 596, 285–290 (2021).

5. Marino, A. et al. Aging and diet alter the protein ubiquitylation landscape in the mouse brain. Nat. Commun. 16, 1–17 (2025).

6. Snyder, N. A. & Silva, G. M. Deubiquitinating enzymes (DUBs): Regulation, homeostasis, and oxidative stress response. J. Biol. Chem. 297, 101077 (2021).

7. Kawaguchi, Y. et al. CAG expansions in a novel gene for Machado-Joseph disease at chromosome 14q32.1. Nat. Genet. 8, 221–228 (1994).

8. Paulson, H. L., Shakkottai, V. G., Clark, H. B. & Orr, H. T. Polyglutamine spinocerebellar ataxias - from genes to potential treatments. Nat. Rev. Neurosci. 18, 613–626 (2017).

9. Leroy, E. et al. The ubiquitin pathway in Parkinson’s disease. Nature 395, 451–452 (1998).

10. Amer-Sarsour, F., Kordonsky, A., Berdichevsky, Y., Prag, G. & Ashkenazi, A. Deubiquitylating enzymes in neuronal health and disease. Cell Death Dis. 12, 120 (2021).

11. Lee, J.-G., Baek, K., Soetandyo, N. & Ye, Y. Reversible inactivation of deubiquitinases by reactive oxygen species in vitro and in cells. Nat. Commun. 4, 1568 (2013).

12. Vozandychova, V. et al. Modified activities of macrophages’ deubiquitinating enzymes after Francisella infection. Front. Immunol. 14, 1252827 (2023).

13. de Jong, A. et al. Ubiquitin-based probes prepared by total synthesis to profile the activity of deubiquitinating enzymes. Chembiochem 13, 2251–2258 (2012).

14. Ekkebus, R. et al. On terminal alkynes that can react with active-site cysteine nucleophiles in proteases. J. Am. Chem. Soc. 135, 2867–2870 (2013).

15. Borodovsky, A. et al. Chemistry-based functional proteomics reveals novel members of the deubiquitinating enzyme family. Chem. Biol. 9, 1149–1159 (2002).

16. Sowa, M. E., Bennett, E. J., Gygi, S. P. & Harper, J. W. Defining the human deubiquitinating enzyme interaction landscape. Cell 138, 389–403 (2009).

17. Ramirez, J. et al. A proteomic approach for systematic mapping of substrates of human deubiquitinating enzymes. Int. J. Mol. Sci. 22, 4851 (2021).

18. Zhang, Y. et al. An RNA-sequencing transcriptome and splicing database of glia, neurons, and vascular cells of the cerebral cortex. J. Neurosci. 34, 11929–11947 (2014).

19. Park, S.-S., Do, H.-A., Park, H.-B., Choi, H.-S. & Baek, K.-H. Deubiquitinating enzyme YOD1 deubiquitinates and destabilizes α-synuclein. Biochem. Biophys. Res. Commun. 645, 124–131 (2023).

20. Tanji, K. et al. YOD1 attenuates neurogenic proteotoxicity through its deubiquitinating activity. Neurobiol. Dis. 112, 14–23 (2018).

21. Zhang, M., Cai, F., Zhang, S., Zhang, S. & Song, W. Overexpression of ubiquitin carboxyl-terminal hydrolase L1 (UCHL1) delays Alzheimer’s progression in vivo. Sci. Rep. 4, 7298 (2014).

22. Mi, Z. & Graham, S. H. Role of UCHL1 in the pathogenesis of neurodegenerative diseases and brain injury. Ageing Res. Rev. 86, 101856 (2023).

23. Day, I. N. M. & Thompson, R. J. UCHL1 (PGP 9.5): Neuronal biomarker and ubiquitin system protein. Prog. Neurobiol. 90, 327–362 (2010).

24. Rott, R. et al. α-Synuclein fate is determined by USP9X-regulated monoubiquitination. Proc. Natl. Acad. Sci. U. S. A. 108, 18666–18671 (2011).

25. Uhlén, M. et al. Proteomics. Tissue-based map of the human proteome. Science 347, 1260419 (2015).

26. Dobson-Stone, C. et al. CYLD is a causative gene for frontotemporal dementia - amyotrophic lateral sclerosis. Brain 143, 783–799 (2020).

27. Di Fraia, D. et al. Altered translation elongation contributes to key hallmarks of aging in the killifish brain. Science 389, eadk3079 (2025).

28. Lushchak, V. I. Free radicals, reactive oxygen species, oxidative stress and its classification. Chem. Biol. Interact. 224, 164–175 (2014).

29. Grintzalis, K. et al. Alterations in thiol redox state and lipid peroxidation in the brain areas of male mice during aging. Adv. Redox Res. 6, 100043 (2022).

30. Ellman, G. L. Tissue sulfhydryl groups. Arch. Biochem. Biophys. 82, 70–77 (1959).

31. Cao, W. et al. Quantifying sulfhydryl oxidation rates using Ellman’s procedure. Phys. Fluids (1994) 37, 017116 (2025).

32. Altun, M. et al. Activity-based chemical proteomics accelerates inhibitor development for deubiquitylating enzymes. Chem. Biol. 18, 1401–1412 (2011).

33. Wang, C. et al. Scalable Production of iPSC-Derived Human Neurons to Identify Tau-Lowering Compounds by High-Content Screening. Stem Cell Reports 9, 1221 (2017).

34. Antico, O. et al. Global ubiquitylation analysis of mitochondria in primary neurons identifies endogenous Parkin targets following activation of PINK1. Sci. Adv. 7, eabj0722 (2021).

35. Ordureau, A. et al. Global landscape and dynamics of Parkin and USP30-dependent ubiquitylomes in iNeurons during mitophagic signaling. Mol. Cell 77, 1124–1142.e10 (2020).

36. Pantazis, C. B. et al. A reference human induced pluripotent stem cell line for large-scale collaborative studies. Cell Stem Cell 29, 1685–1702.e22 (2022).

37. Cowell, I. G., Ling, E. M., Swan, R. L., Brooks, M. L. W. & Austin, C. A. The deubiquitinating enzyme inhibitor PR-619 is a potent DNA topoisomerase II poison. Mol. Pharmacol. 96, 562–572 (2019).

38. Fulzele, A. & Bennett, E. J. Ubiquitin diGLY proteomics as an approach to identify and quantify the ubiquitin-modified proteome. Methods Mol. Biol. 1844, 363–384 (2018).

39. Kim, W. et al. Systematic and quantitative assessment of the ubiquitin-modified proteome. Mol. Cell 44, 325–340 (2011).

40. Umansky, C. et al. Endogenous formaldehyde scavenges cellular glutathione resulting in redox disruption and cytotoxicity. Nat. Commun. 13, 745 (2022).

41. Yao, T. et al. Distinct modes of regulation of the Uch37 deubiquitinating enzyme in the proteasome and in the Ino80 chromatin-remodeling complex. Mol. Cell 31, 909–917 (2008).

42. Lee, B.-H. et al. Enhancement of proteasome activity by a small-molecule inhibitor of USP14. Nature 467, 179–184 (2010).

43. Matilainen, O., Arpalahti, L., Rantanen, V., Hautaniemi, S. & Holmberg, C. I. Insulin/IGF-1 signaling regulates proteasome activity through the deubiquitinating enzyme UBH-4. Cell Rep. 3, 1980–1995 (2013).

44. Kim, H. T. & Goldberg, A. L. The deubiquitinating enzyme Usp14 allosterically inhibits multiple proteasomal activities and ubiquitin-independent proteolysis. J. Biol. Chem. 292, 9830–9839 (2017).

45. Seiberlich, V., Goldbaum, O., Zhukareva, V. & Richter-Landsberg, C. The small molecule inhibitor PR-619 of deubiquitinating enzymes affects the microtubule network and causes protein aggregate formation in neural cells: implications for neurodegenerative diseases. Biochim. Biophys. Acta 1823, 2057–2068 (2012).

46. Giustarini, D., Milzani, A., Dalle-Donne, I., Tsikas, D. & Rossi, R. N-Acetylcysteine ethyl ester (NACET): a novel lipophilic cell-permeable cysteine derivative with an unusual pharmacokinetic feature and remarkable antioxidant potential. Biochem. Pharmacol. 84, 1522–1533 (2012).

47. Tosi, G. M. et al. Superior properties of N-acetylcysteine ethyl ester over N-acetyl cysteine to prevent retinal pigment epithelial cells oxidative damage. Int. J. Mol. Sci. 22, 600 (2021).

48. Vilchez, D., Saez, I. & Dillin, A. The role of protein clearance mechanisms in organismal ageing and age-related diseases. Nat. Commun. 5, 5659 (2014).

49. Nago, N., Murata, S., Tanaka, K. & Tanahashi, N. Changes in brain proteasome dynamics associated with aging. Genes Cells 29, 438–445 (2024).

50. Davidson, K. & Pickering, A. M. The proteasome: A key modulator of nervous system function, brain aging, and neurodegenerative disease. Front. Cell Dev. Biol. 11, 1124907 (2023).

51. Munkácsy, E. et al. Neuronal-specific proteasome augmentation via Prosβ5 overexpression extends lifespan and reduces age-related cognitive decline. Aging Cell 18, e13005 (2019).

52. Vernace, V. A., Arnaud, L., Schmidt-Glenewinkel, T. & Figueiredo-Pereira, M. E. Aging perturbs 26S proteasome assembly in Drosophila melanogaster. FASEB J. 21, 2672–2682 (2007).

53. Weishäupl, D. et al. Physiological and pathophysiological characteristics of ataxin-3 isoforms. J. Biol. Chem. 294, 644–661 (2019).

54. Hernández-Carralero, E., Quinet, G. & Freire, R. ATXN3: a multifunctional protein involved in the polyglutamine disease spinocerebellar ataxia type 3. Expert Rev. Mol. Med. 26, e19 (2024).

55. Du, F., Yu, Q., Kanaan, N. M. & Yan, S. S. Mitochondrial oxidative stress contributes to the pathological aggregation and accumulation of tau oligomers in Alzheimer’s disease. Hum. Mol. Genet. 31, 2498–2507 (2022).

56. Zong, Y. et al. Mitochondrial dysfunction: mechanisms and advances in therapy. Signal Transduct. Target. Ther. 9, 124 (2024).

57. Bin, P., Huang, R. & Zhou, X. Oxidation resistance of the sulfur amino acids: Methionine and cysteine. Biomed Res. Int. 2017, 9584932 (2017).

58. Gusarov, I. et al. Dietary thiols accelerate aging of C. elegans. Nat. Commun. 12, 4336 (2021).

59. Bulteau, A.-L., Szweda, L. I. & Friguet, B. Mitochondrial protein oxidation and degradation in response to oxidative stress and aging. Exp. Gerontol. 41, 653–657 (2006).

60. Cotto-Rios, X. M., Békés, M., Chapman, J., Ueberheide, B. & Huang, T. T. Deubiquitinases as a signaling target of oxidative stress. Cell Rep. 2, 1475–1484 (2012).

61. Rizza, S. & Filomeni, G. Denitrosylate and live longer: how ADH5/GSNOR links mitophagy to aging. Autophagy 14, 1285–1287 (2018).

62. Kou, Y., Zhao, H., Cui, D., Han, H. & Tong, Z. Formaldehyde toxicity in age-related neurological dementia. Ageing Res. Rev. 73, 101512 (2022).

63. Loeffler, D. A., Klaver, A. C., Coffey, M. P., Aasly, J. O. & LeWitt, P. A. Age-related decrease in heat shock 70-kDa protein 8 in cerebrospinal fluid is associated with increased oxidative stress. Front. Aging Neurosci. 8, 178 (2016).

64. Sirtori, R., Riva, C., Ferrarese, C. & Sala, G. HSPA8 knock-down induces the accumulation of neurodegenerative disorder-associated proteins. Neurosci. Lett. 736, 135272 (2020).

65. Panda, P., Sarohi, V., Basak, T. & Kasturi, P. Elucidation of site-specific ubiquitination on chaperones in response to mutant huntingtin. Cell. Mol. Neurobiol. 44, 3 (2023).

66. Griffith, A. A. & Holmes, W. Fine tuning: Effects of post-translational modification on Hsp70 chaperones. Int. J. Mol. Sci. 20, 4207 (2019).

67. Shan, S. et al. Marine algae-derived oligosaccharide via protein crotonylation of key targeting for management of type 2 diabetes mellitus in the elderly. Pharmacol. Res. 205, 107257 (2024).

68. Shriya, S., Paul, R., Singh, N., Afza, F. & Jain, B. P. Bioinformatics analysis and alternative polyadenylation in Heat Shock Proteins 70 (HSP70) family members. Int. J. Physiol. Pathophysiol. Pharmacol. 16, 138–151 (2024).

69. Ainatzi, S. et al. Ca2+-triggered (de)ubiquitination events in synapses. Mol. Cell. Proteomics 24, 100946 (2025).

70. Chen, H., Polo, S., Di Fiore, P. P. & De Camilli, P. V. Rapid Ca2+-dependent decrease of protein ubiquitination at synapses. Proc. Natl. Acad. Sci. U. S. A. 100, 14908–14913 (2003).

71. Sheng, Y. et al. Molecular recognition of p53 and MDM2 by USP7/HAUSP. Nat. Struct. Mol. Biol. 13, 285–291 (2006).

72. Stevenson, L. F. et al. The deubiquitinating enzyme USP2a regulates the p53 pathway by targeting Mdm2. EMBO J. 26, 976–986 (2007).

73. Mulder, M. P. C. et al. A cascading activity-based probe sequentially targets E1-E2-E3 ubiquitin enzymes. Nat. Chem. Biol. 12, 523–530 (2016).

74. Rao, N. R., Upadhyay, A. & Savas, J. N. Derailed protein turnover in the aging mammalian brain. Mol. Syst. Biol. 20, 120–139 (2024).

75. Srinivasan, V. et al. USP14 is crucial for proteostasis regulation and α-synuclein degradation in human SH-SY5Y dopaminergic cells. Heliyon 11, e42031 (2025).

76. Kors, S., Geijtenbeek, K., Reits, E. & Schipper-Krom, S. Regulation of proteasome activity by (post-)transcriptional mechanisms. Front. Mol. Biosci. 6, 48 (2019).

77. Besche, H. C. et al. Autoubiquitination of the 26S proteasome on Rpn13 regulates breakdown of ubiquitin conjugates. EMBO J. 33, 1159–1176 (2014).

78. Isasa, M. et al. Monoubiquitination of RPN10 regulates substrate recruitment to the proteasome. Mol. Cell 38, 733–745 (2010).

79. Reeg, S., Castro, J. P., Hugo, M. & Grune, T. Accumulation of polyubiquitinated proteins: A consequence of early inactivation of the 26S proteasome. Free Radic. Biol. Med. 160, 293–302 (2020).

80. Ishii, T., Sakurai, T., Usami, H. & Uchida, K. Oxidative modification of proteasome: identification of an oxidation-sensitive subunit in 26 S proteasome. Biochemistry 44, 13893–13901 (2005).

81. Hugo, M. et al. Early cysteine-dependent inactivation of 26S proteasomes does not involve particle disassembly. Redox Biol. 16, 123–128 (2018).

82. Livnat-Levanon, N. et al. Reversible 26S proteasome disassembly upon mitochondrial stress. Cell Rep. 7, 1371–1380 (2014).

83. Roux, A. E. et al. Individual cell types in C. elegans age differently and activate distinct cell-protective responses. Cell Rep. 42, 112902 (2023).

84. Chien, J.-F. et al. Cell-type-specific effects of age and sex on human cortical neurons. Neuron 112, 2524–2539.e5 (2024).

85. Khawaja, R. R. et al. Sex-specific and cell-type-specific changes in chaperone-mediated autophagy across tissues during aging. *Nat*. Aging 5, 691–708 (2025).

86. Peth, A., Kukushkin, N., Bossé, M. & Goldberg, A. L. Ubiquitinated proteins activate the proteasomal ATPases by binding to Usp14 or Uch37 homologs. J. Biol. Chem. 288, 7781–7790 (2013).

87. Bingol, B. et al. The mitochondrial deubiquitinase USP30 opposes parkin-mediated mitophagy. Nature 510, 370–375 (2014).

88. Fang, T.-S. Z. et al. Knockout or inhibition of USP30 protects dopaminergic neurons in a Parkinson’s disease mouse model. Nat. Commun. 14, 7295 (2023).

89. Zhang, X.-W. et al. Neuroinflammation inhibition by small-molecule targeting USP7 noncatalytic domain for neurodegenerative disease therapy. Sci. Adv. 8, eabo0789 (2022).

90. VerPlank, J. J. S., Lokireddy, S., Feltri, M. L., Goldberg, A. L. & Wrabetz, L. Impairment of protein degradation and proteasome function in hereditary neuropathies. Glia 66, 379–395 (2018).

91. Kim, E. et al. Dual function of USP14 deubiquitinase in cellular proteasomal activity and autophagic flux. Cell Rep. 24, 732–743 (2018).

92. Qin, B., Chen, X., Wang, F. & Wang, Y. DUBs in Alzheimer’s disease: mechanisms and therapeutic implications. Cell Death Discov. 10, 475 (2024).

93. Naumann, U., Brazzell, J. L., Crim, M. J. & Hoppe, B. Comprehensive colony health management and emerging pathogens of the annual killifish species Nothobranchius furzeri. J. Am. Assoc. Lab. Anim. Sci. 63, 20–33 (2024).

94. Cirri, E. et al. Optimized automated workflow for BioID improves reproducibility and identification of protein-protein interactions. J. Proteome Res. 23, 4359–4368 (2024).

95. Buczak, K. et al. Spatially resolved analysis of FFPE tissue proteomes by quantitative mass spectrometry. Nat. Protoc. 15, 2956–2979 (2020).

96. Wu, T. et al. clusterProfiler 4.0: A universal enrichment tool for interpreting omics data. Innovation (Camb.) 2, 100141 (2021).

